# Ferroptosis governs lymphatic vessel growth and regression

**DOI:** 10.1101/2025.11.18.689033

**Authors:** Maria-Kyriaki Drekolia, Daiyu Wang, Yunyun Chen, Yali Du, Janina Wittig, Ran Xu, Danny Schilling, Blerina Aliraj, Janina Mettner, Lavanya Gupta, Cong Xu, Stefan Günther, Ronay Cetin, Yves Matthess, Ingrid Fleming, Manuel Kaulich, Gergana Dobreva, Andreas Weigert, Tobias P. Dick, Ivan Dikic, Alexandra Stolz, Karina Yaniv, Andreas Linkermann, Jiong Hu, Sofia-Iris Bibli

**Author notes:** These authors contributed equally, listed in alphabetical order (Maria-Kyriaki Drekolia, Daiyu Wang). **Correspondence:** <u>Sofia-Iris Bibli, PhD:</u> Department of Vascular Dysfunction, European Center for Angioscience (ECAS), Medical Faculty Mannheim, Heidelberg University, Mannheim, Ludolf-Krehl-Straße 13-17, 68167 Mannheim, Germany.

## Abstract

Whether, when, and how lymphatic vessels undergo cell death remains poorly understood. Here we identify ferroptosis as a physiological, cell-intrinsic regulator of the lymphatic endothelial cell survival during development and following injury, in stark contrast to the resilient organotypic blood endothelial cells. The lymphatic susceptibility to ferroptosis stems from tampered cystine/ hydropersulfide metabolism, alongside reduced glutathione availability triggered by an SH3RF3 E3 ligase mediated GPX4 degradation, and enhanced integration of polyunsaturated fatty acid enriched membrane phospholipids. Inducing ferroptosis genetically or pharmacologically elevated lymphatic lipid peroxidation, halted embryonic lymphangiogenesis and prevented post-injury lymphatic overgrowth while simultaneously shaped immune responses. Conversely, ferroptosis inhibition through saturated fatty acid supplementation led to pathological lymphatic hyperplasia. Targeting lymphatic ferroptotic mechanisms holds promise against pathological lymphatic growth in response to injury.

## Main Text

The lymphatic system plays a vital role in maintaining tissue fluid balance, facilitating immune surveillance, and clearing metabolic waste (*1, 2*). The discovery of lymphatic endothelial cell-specific markers has significantly advanced our understanding of lymphatic vessel distribution and function in organs once thought to lack them, including the brain, eye, and bone (*3–11*). Lymphatic endothelial cells arise from a specialized niche of angioblasts via trans-differentiation and respond to mechanical cues to proliferate (*12–14*). This process initiates lymphangiogenesis -the formation of the lymphatic network from preexisting vasculature (*15*)-around embryonic day (E) 9.5, driven by the transcription factor PROX1 and further regulated by SOX18 and COUP-TFII (*16, 17*). VEGF-C signaling then promotes the budding of lymphatic progenitors from the cardinal vein, establishing the primitive lymph sacs (*18*). More recently, BMP2 (*19*) and the RAF1/MEK/ERK signalling cascade (*20*) have also been implicated in lymphatic fate specification. While lymphangiogenesis is highly active during embryonic development, in adults it is typically restricted to specific conditions, such as the female reproductive cycle, wound healing, and pathological contexts such as tumors, inflammation (*21*), or genotoxic stress (*4*). Inadequate lymphatic expansion in these settings can contribute to diseases like lymphedema (*1*).

Despite significant progress in understanding lymphangiogenic signaling (*22*), the mechanisms governing lymphatic endothelial cells turnover and vessel regression remain poorly defined. During both development (*23*) and tissue repair (*4, 24*), blood and lymphatic vessels form networks, that through remodeling and maturation, shape an efficient, hierarchical vasculature. While the endothelial cells of blood vessels predominantly regress through well-characterized apoptotic pathways (*25–34*), it is still unclear whether lymphatic vessels rely on similar or distinct mechanisms of programmed cell death. This gap in knowledge is striking, given the highly adaptive and dynamic nature of the lymphatic network. Defining how lymphatic vessels are maintained, remodeled, or eliminated is essential for understanding tissue homeostasis and disease. In this study, we identify physiological ferroptosis as a central and tightly regulated mechanism for lymphatic vessel remodeling. Through tracking both developmental and injury-induced lymphatic growth, we delineate distinct, lymphatic-specific cell death pathways absent in corresponding blood vessels.

## Results

### Lipid peroxidation accompanies lymphatic endothelial cell growth and regression

To identify potential programmed cell death mechanisms operative during lymphatic and blood vessel development, we examined murine embryonic skin, where both vascular networks coexist (**Fig. 1A**). During the developmental stage (E13.5-E18.5) both embryonic blood (ERG1^+^, CD31^+^, PROX1^-^) and lymphatic endothelial cells (PROX1^+^, NRP2^+^) showed no change in the phosphorylation of Mixed lineage kinase domain-like protein (MLKL) at S345, a marker of necroptosis (**fig. S1A, S1B**), whilst only blood endothelial cells, previously reported to initiate apoptosis during their maturation and regression phase (*25–34*), exhibited increased cleaved caspase 3/7 signals at E17.5 (**fig. S1C, S1D**). In contrast, the lipid peroxidation adduct 4-hydroxynonenal (4HNE), a marker of ferroptosis, was found elevated in a subset of growing lymphatics at E14.5 and E15.5 (**Fig. 1B, fig. S1E**), while embryonic blood endothelial cells remained negative during their growth and maturation phase (**Fig. 1C**). Electron microscopy revealed mitochondrial structural damage with loss of cristae structure, but no differences in the nuclear membrane integrity, specifically in lymphatic endothelial cells at E15.5, while blood endothelial cells mitochondria morphology remained intact at this time point (**Fig. 1D**). This coincided with increased lymphatic mitochondria lipid peroxidation signals (**fig. S1F**), confirming that excessive lipid peroxidation is a hallmark of prenatal lymphatic but not blood vessel development. To extend these findings, we next analyzed postnatal corneal lymphatic vessels, in which lymphatic regression was observed between postnatal (p) days 16 to 22 (**Fig. 1E**). Similar to the situation in embryonic skin, a subset of proliferating (EdU^+^) corneal lymphatic endothelial cells contained high levels of 4HNE that were even higher in non-proliferating (EdU^-^) cells, peaking at p16 (during the regression phase) and returning to baseline at p22 (**Fig. 1F, fig. S1G**). Notably, corneal lymphatic endothelial cells, showed no signs of phosphatidylserine externalization or membrane rupture (*35*) as indicated by the absence of Annexin V signals during development and regression, suggesting that programmed cell death signals that disrupt membrane integrity (*36–38*) are potentially not a characteristic feature of post natal lymphatic vessel growth and regression. Mitochondria lipid peroxidation followed the trajectory of 4HNE, starting at p16 and remaining elevated until p20, when the corneal lymphatic regression was completed (**fig. S1H**). To further assess whether increased lipid peroxidation actually results in ferroptosis and actively regulates lymphatic regression, we administered the ferroptosis inhibitor, ferrostatin 1, at p14; the onset of the regression phase. Inhibiting ferroptosis reduced mitochondrial lipid peroxidation (**fig. S1I**), decreased 4HNE signals and led to lymphatic hyperplasia at p16 (**Fig. 1G**). Taken together these findings suggest that ferroptosis is a physiological surveillance mechanism required to fine-tune prenatal lymphatic expansion and postnatal regression.

**Fig. 1.**
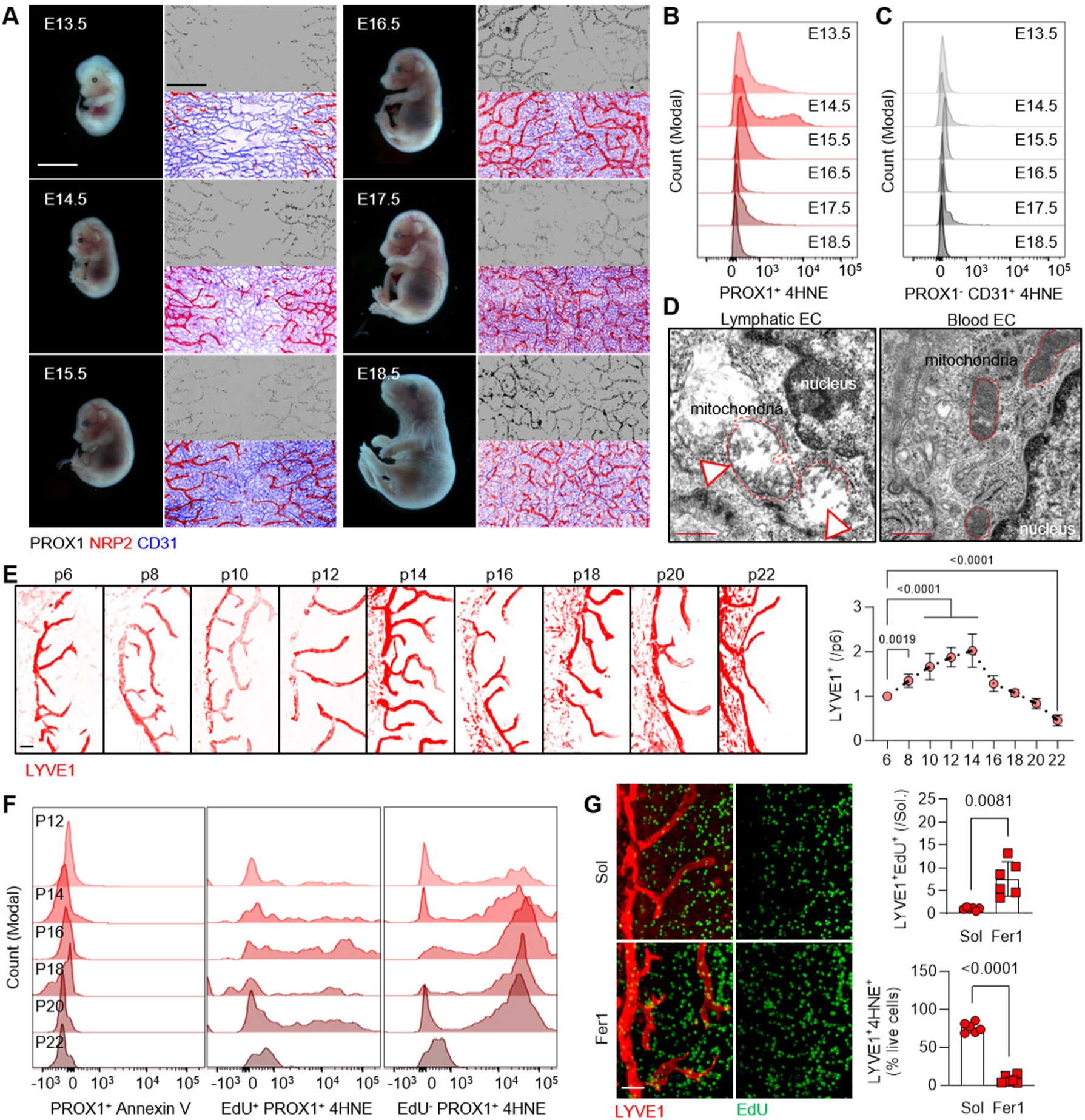
Ferroptosis governs lymphatic endothelial cell development. (**A**) Stereomicrographic images of murine embryos from embryonic day (E) E13.5 to E18.5 alongside immunofluorescent images of CD31 (blue) blood, PROX1 (black) and NRP2 (red) dermal lymphatic endothelial cells. Scale bars = 1 mm and 200 μm. Similar results were obtained from 4 additional embryos per time point. (**B-C**) Intensities of 4-hydroxynonenal (4HNE) in PROX1^+^ dermal lymphatic (B) and PROX1^-^CD31^+^ blood (C) endothelial cells isolated from samples as in panel (A) following flow cytometry analysis. n= 8 embryos/ group. (**D**) Transmission electron microscopy of the dermal lymphatic and blood endothelial cells from embryo skin at E15.5. Arrows: mitochondrial morphology; dashed red lines: mitochondria. Scale bars = 1 μm. (**E**) Immunostaining of LYVE1 (red) lymphatic endothelial cells from murine cornea preparations at postnatal (p) days 6 to 22 and respective quantification. Scale bars = 100 μm, n=5-7/group. Kolmogorov-Smirnov normality test, One way Anova (Dunnett’s multiple comparisons test). (**F**) Intensities of Annexin V or 4HNE from the PROX1^+^ population in proliferating (EdU^+^) or quiescent (EdU^-^) lymphatic endothelial cells isolated from murine postnatal cornea following flow cytometry analysis. n=pool of 8 corneas/group. (**G**) Immunostaining of LYVE1 (red) lymphatic endothelial cells and EdU (green) proliferating cells on p16 murine cornea after receiving treatment with solvent (Sol) or 50 μmol/L Ferrostatin 1 (Fer1) from p14 and every 12 hours, and respective quantification of LYVE1^+^ EdU^+^ and percentage of LYVE1^+^ 4HNE^+^ live cells. n=6 corneas/ group. Scale bars = 100 μm. Kolmogorov-Smirnov normality test, paired Student’s *t* test.

### Cystine import and glutathione generation shield the lymphatic endothelium from ferroptosis

To determine how lymphatic endothelial cells are uniquely sensitive to lipid peroxidation and ferroptotic cell death, we investigated differences in the molecular mechanisms governing ferroptosis (*39*). The key characteristic of ferroptotic cell death is the iron-dependent phospholipid peroxidation through Fenton and Haber-Weiss reactions (*40*) (**fig. S2A**); distinguishing it from apoptosis, necrosis and autophagy-dependent cell death. By comparing human blood and lymphatic endothelial cells we indeed observed increased basal levels of the labile Fe^2+^ within the lymphatic endothelial cell (**fig. S2B**), which were sensitive to the iron chelator deferoxamine (DFO) (**fig. S2B**), consistent with previous observations (*40*). The accumulation of Fe^2+^ could be the consequence of mild transcriptional differences in the ferritin transporter (TFRC), or potentially the ferritin heavy chain 1 (FTH1); but rather not the differential expression of the endosomal metalloreductase STEAP3, which reduces Fe^3+^(**fig. S2C**). The increased labile iron in the lymphatic endothelium compared to the organotypic blood endothelium correlated with increased basal levels of lipid peroxidation, as indicated by increased C11-Bodipy oxidation (**fig. S2D**), mitochondria reactive oxygen species (**fig. S2E**) and lipid peroxidation signals (**fig. S2F**).

To counteract iron-dependent lethal lipid peroxidation, cells employ multiple ferroptosis-suppressing pathways primarily including the mevalonate β-Hydroxy β-methylglutaryl-(HMG) CoA pathway, the ferroptosis suppressor protein 1 (FSP1)/CoQ10/ubiquinone antioxidant defense pathway, the biopterin dihydrofolate reductase system as well as the cystine/glutathione metabolic pathway (**fig. S3A**) (*39*). It was possible to exclude the mevalonate pathway as a major regulator of lymphatic cell death by comparing the transcriptome of blood vs. lymphatic juvenile and adult skin endothelial cells. Such analysis revealed no significant differences in the enzymes responsible for the mevalonate to cholesterol biosynthesis, except of a mild reduction in the transcriptional levels of the ferroptosis suppressor sterol C5 desaturase (SC5D) (**fig. S3B**). Targeted metabolomic studies also failed to detect any significant differences in cholesterol levels or its intermediate product 7-dehydrocholesterol (7-DHC) between blood and lymphatic endothelial cells (**fig. S3C**). Mevalonate-derived vitamin K has also been implicated in protection against ferroptosis (*41*), however, no direct differences in its levels were detected in the two cell types (**fig. S3D**).

The mevalonate pathway acts also as a key carbon source for the synthesis of CoQ10 (*42*) which is a major suppressor of ferroptosis (*43, 44*) and the main effector of the FSP1 pathway (*45*). However, we found ferroptosis in the lymphatic endothelial cells to be independent of the mevalonate/cholesterol/FSP1 axis. In particular, while the acetyl CoA precursor 3-hydroxy-3-methylglutaramate (*46*), was elevated in the blood endothelial cells (**fig. S3E**), total CoQ10 levels were similar between the two cell types (**fig. S3F**). FSP1 but not dihydroorotate dehydrogenase (DHODH) transcriptional levels were found reduced in the lymphatic endothelium (**fig. S3G**) and NAD^+^ levels (**fig. S3H**) were significantly higher and a-tocopherol levels lower within the blood endothelium (**fig. S3I**); yet, inhibiting FSP1 failed to induce basal ferroptotic death in either cell types (**fig. S2J**).

The canonical ferroptosis process entails uptake of cystine via the system xc-cystine/glutamate antiporter (SLC7A11/ SLC3A2 subunits), followed by thioredoxin reductase 1 (TXNRD1)-dependent reduction of cystine to cysteine (*39*). Once taken up, cystine/cysteine support the biosynthesis of glutathione (GSH) which opposes ferroptosis by contributing to glutathione peroxidase (GPx) activity (**Fig. 2A**) (*39*). Although, only dermal adult lymphatic endothelial cells showed reduced transcription of the SLC7A11 (**fig. S4A**), this was independent from an OPA1 mediated activating transcription factor 4 (ATF4) mechanism (**fig. S4B**) as recently described (*47*). SLC7A11 protein levels were significantly lower in dermal lymphatic endothelial cells (**Fig. 2B**) which resulted in decreased imported 13C-cystine compared to the dermal blood endothelial cells (**Fig. 2C**). Interestingly, lymphatic endothelial cells relied heavily (∼80%) on carbons from cystine to generate GSH, in contrast to blood endothelial cells (**Fig. 2D**), although the overall levels of lymphatic endothelial cell GSH were significantly lower (**Fig. 2E**). These data suggest a strong dependence of lymphatic endothelial cells on the cystine-GSH axis. Indeed, lymphatic endothelial cells survival and proliferation exhibited stronger dependency on increasing concentrations of cystine compared to blood endothelial cells (**Fig. 2F, Supplemental Videos 1A-1B**). Inhibition of cystine import, by IKE; a selective inhibitor of the SLC7A11 antiporter, resulted in a concentration dependent death of the lymphatic endothelial cells with the characteristic morphological ferroptotic features (**Fig. 2G, Supplemental Videos 1C-1D**). In contrast, inhibition of the SLC7A11 antiporter reduced proliferation of blood endothelial cells, without initiating cell death (**Fig. 2G**). Lymphatic endothelial cell death initiated by cystine removal or inhibition of cystine import, could be rescued only by the addition of N-acetyl-cysteine, the ferroptosis inhibitor ferrostatin 1, the iron chelator deferoxamine or the lipophilic antioxidant (soluble Vitamin E analogue) Trolox (**Fig. 2H**). Inhibitors of apoptosis, necroptosis and autophagy failed to maintain lymphatic endothelial cell survival (**Fig. 2H**). Notably, ferroptosis could not be induced in blood endothelial cells by either cystine removal or SLC7A11 inhibition (**Fig. 2F-2H**). Silencing SLC7A11 (**fig. S4C**) in lymphatic endothelial cells resulted in halted proliferation and increased signals of the cell death indicator, Sytox (**fig. S4D-S4E**). *In vivo* inducible deletion of lymphatic SLC7A11, to inhibit import of cystine, during the developmental stage of lymphatics i.e. E9.5, resulted in embryos with excessive edema at E15.5, impaired lymphatic network development and fewer filopodia (**Fig. 2I**), as well as increased accumulation of 4HNE (**Fig. 2J**). Taken together these data suggest that a cystine dependent endothelial ferroptotic death mechanism is unique for the lymphatic endothelium.

**Fig. 2.**
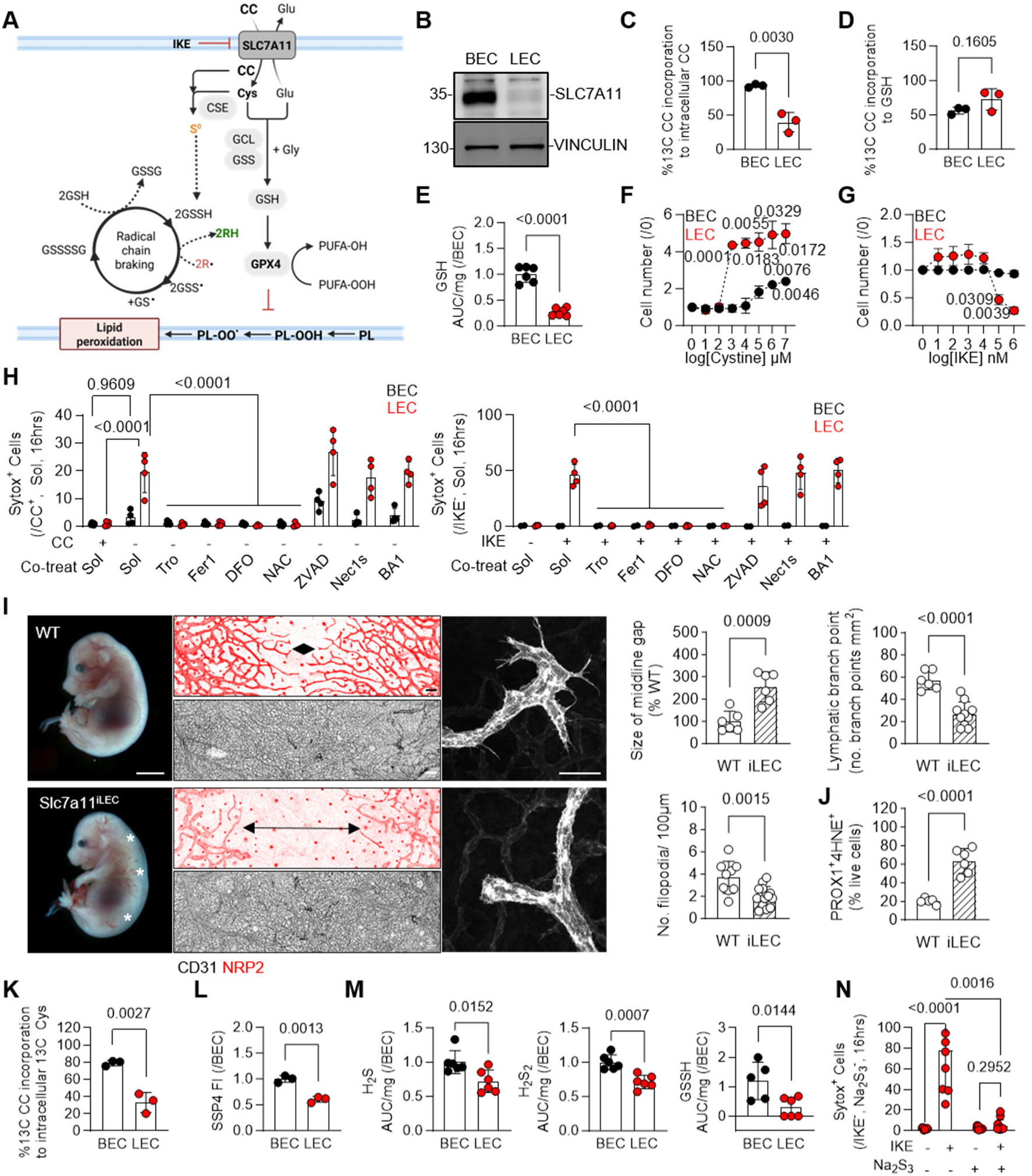
Reduced cystine import and catabolism induces ferroptosis and compromises lymphatic vessel growth. (**A**) Schematic showing the pathway of cystine import and flux to persulfides and glutathione in mammalian cells. SLC7A11: Solute carrier family 7 member 11; CSE: Cystathionine gamma-Lyase; GCL: glutamate-cysteine ligase; GSS: glutathione synthetase; GPX4: glutathione peroxidase 4; CC: Cystine; Glu: Glutamate; Cys: Cysteine; Gly: Glycine; GSH: Glutathione; GSSG: Glutathione disulfide; S^0^: Elemental sulfur; GSSSSG: Glutathione polysulfide; PUFA: Polyunsaturated fatty acid; IKE: Imidazole ketone erastin. (**B**) Representative immunoblot of SLC7A11 and VINCULIN in human lymphatic (LEC) and blood (BEC) endothelial cells. Similar data were obtained from n=3 additional batches. (**C**) Percentage of 13C Cystine (CC) import in human lymphatic (LEC) and blood (BEC) endothelial cells. n=3/ group in 3 independent experiments. Sharipo-Wilk normality test. Unpaired Student‘s t test. (**D**) Percentage of 13C Cystine (CC) incorporation to GSH in samples as in panel (C). n=3/ group. Sharipo-Wilk normality test. Unpaired Student‘s t test. (**E**) Levels of GSH in human lymphatic (LEC) and blood (BEC) endothelial cells. n=6/ group. Kolmogorov-Smirnov normality test. Unpaired Student‘s t test. (**F**) Relative cell number of human lymphatic (LEC) and blood (BEC) endothelial cells in response to increasing concentrations of cystine at 12 hours. n=3/ group in 3 independent experiments. Two way Anova, Dunnet’s multiple comparisons test. (**G**) Relative cell number of human lymphatic (LEC) and blood (BEC) endothelial cells in response to increasing concentrations of the selective SLC7A11 inhibitor IKE (imidazole ketone erastin) at 12 hours. n=3/ group in 3 independent experiments. Two way Anova, Dunnet’s multiple comparisons test. (**H**) Relative Sytox positive human dermal lymphatic (LEC) and blood (BEC) endothelial cells, cultured in the presence or absence of 100 µmol/L cystine (CC) (right panel) or treated with Solvent (Sol) or 10 µmol/L IKE (left panel) and simultaneously exposed to Trolox (Tro, 100μmol/L), Ferrostatin 1 (Fer1, 10μmol/L), Deforoxamine (DFO, 100 μmol/L), N-acetyl-L-cysteine (NAC, 1 mmol/L), Z-VAD-FMK Caspase inhibitor (ZVAD, 50μmol/L), Necrostatin (Nec1s, 10 μmol/L) or Bafilomycin 1 (BA1, 1 nmol/L) for 16 hours respectively. n=4/ group (3 technical replicates) in 3 independent experiments. Two way Anova, Dunnet’s multiple comparisons test. (**I**) Stereomicrographic images of E15.5 embryos and immunofluorescent images of CD31 (black) blood and NRP2 (red) dermal lymphatic endothelial cells in Prox1 Cre^-^ (WT) or Cre^+^ (Slc7a11^iLEC^) mice treated with tamoxifen from E10.5-E12.5 to induce genetic deletion of lymphatic endothelial cell *Slc7a11*. Right panel shows NRP2 (gray) lymphatic filopodia in high magnification. Relative quantification of size of middline gap as % to WT, number of filopodia and lymphatic branch points in the same embryos. Asterisks: murine edema. Black arrows: indicator of middline gap. White arrows: filopodia. Scale bars = 1 mm, 200 μm and 50 μm respectively. n=6 embryos/ group from 2 independent births. Kolmogorov-Smirnov normality test. Unpaired Student‘s t test. (**J**) Percentage of PROX1^+^ 4HNE^+^ live cells in samples as in panel (I). n=6 embryos/ group. Kolmogorov-Smirnov normality test. Unpaired Student‘s t test. (**K**) Percentage of 13C Cystine (CC) incorporation to cysteine (Cys) in human lymphatic (LEC) and blood (BEC) endothelial cells. n=3/ group (2 technical replicates) in 3 independent experiments. Sharipo-Wilk normality test. Unpaired Student‘s t test. (**L**) Sulfane sulfur species levels detected by the relative fluorescent SSP4 intensity (FI) in human lymphatic (LEC) and blood (BEC) endothelial cells exposed to solvent or the iron chelator deferoxamine (DFO, 100 μmol/L, 8 hours) following flow cytometry analysis. n=3/ group. Sharipo-Wilk normality test. Unpaired Student‘s t test. (**M**) Levels of H_2_S, H_2_S_2_ and GSSH in human lymphatic (LEC) and blood (BEC) endothelial cells. n=5-6/ group. H_2_S levels: Mann-Whitney test, H_2_S_2_: Shapiro-Wilk normality test, unpaired Student‘s t test, GSSH: Kolmogorov-Smirnov normality test, unpaired Student‘s t test. (**N**) Relative Sytox positive human dermal lymphatic endothelial cells treated with 10 µmol/L IKE for 16 hours after pre-treatment with the persulfide donor Na_2_S_3_ (10 µmol/L, 24hrs). n=6-7/ group. Two way Anova, Fischer’s multiple comparison test.

Intracellular cystine can also support ferroptosis resistance through enzymatic generation of sulfane sulfur species (*48*), thus we next investigated the possibility that the lack of hydropersulfides render lymphatic endothelial cells sensitive to ferroptosis. Consistent with the reduced cystine uptake, lymphatic endothelial cells contained lower levels of intracellular cysteine (**Fig. 2K**) and sulfane sulfur levels (detected by SSP4), compared to blood endothelial cells (**Fig. 2L**). While blood endothelial cells have the capacity to generate endogenous persulfides through the activity of cystathionine gamma lyase (CSE) (*49*), which are important to maintain multiple blood vessel functions (*50–52*), the role of CSE in lymphatic endothelial cells is unknown. While on a transcriptional level no differences were detected for the CTH gene which encodes the CSE protein (**fig. S4F**), a trend of lower protein expression in the human dermal lymphatic endothelial cells was observed (**fig. S4G**). This difference was even more striking in lymphatic endothelial cells isolated form the murine embryo skin at E 15.5, in which CSE was not highly expressed (**fig. S4H**); results that could potentially explain the low levels of lymphatic endothelial cell persulfides measured *in vivo* (**fig. S4I**). Reduced S^0^ levels within the lymphatic endothelium, were paralleled by reduced generation of hydrogen sulfide, polysulfides and glutathione hydropersulfides (**Fig. 2M**). Addition of a persulfide donor rendered the lymphatic endothelial cell resistant to IKE mediated cell death (**Fig. 2N**). However, deletion of CSE in the blood endothelial cells, to inhibit one persulfide producing pathway (*49, 51*) did not affect significantly their response to IKE mediated cell death (**fig. S4J**); data indicating that additional protective mechanisms against ferroptosis are utilized by the blood endothelium. In summary, our findings reveal an unrecognized; angiodiverse; ferroptotic sensitivity, with lymphatic unlike blood endothelial cells depending on cystine import and glutathione bioavailability for survival.

### GPX4 deletion induces lymphatic death and potentiates lymphatic regression

In line with the cystine dependent sensitivity of the lymphatic vasculature to ferroptosis; pharmacological inhibition of GPX4, confirmed that only the lymphatic endothelium is sensitive to ferroptotic cell death signals (**Fig. 3A-3B, Supplemental Videos 2A-2D**). The latter response was antagonized by the addition of the ferroptosis inhibitor ferrostatin 1, the iron chelator deferoxamine or the lipophilic antioxidant Trolox (**Fig. 3C**). Blood endothelial cells did not undergo GPX4 mediated ferroptosis. Silencing GPX4 (**fig. S5A**) in the lymphatic endothelial cells reduced lymphatic growth and increased cell death (**fig. S5B-S5C**). The PROX1 dependent inducible genetic deletion of *Gpx4* prior to the development of the dermal murine lymphatic system i.e. E9.5, resulted in embryos with edema at E15.5 and a simultaneous reduction in lymphatic growth, branching points and number of filopodia (**Fig. 3D**). Lymphatic endothelial cells in GPX4 depleted littermates, showed increased 4HNE (**Fig. 3E**) and disrupted mitochondrial structure (**Fig. 3F**), pointing to a ferroptosis mediated death mechanism. Intragastrial treatment of the pregnant mothers with ferrostatin 1 reversed completely the phenotypes, re-establishing lymphatic growth at E15.5 and reducing lymphatic 4HNE levels (**Fig. 3D-3E**). In contrast to the consequences of GPX4 deletion in the lymphatic endothelium, the Cdh5-Cre driven inducible deletion of *Gpx4* at E9.5 did not have any impact on blood endothelial cell growth in the murine skin at E15.5 and did not initiate apoptotic (**fig. S5D**) or lipid peroxidation signals (**fig. S5E**). This approach did however elicit mild edema and a reduction in lymphangiognenesis largely attributable to the fact that lymphatic endothelial cells also express Cdh5 (**fig. S5F**). The inducible deletion of *Gpx4* from E14.5-E15.5 resulted in less dense and mature lymphatic vasculature at E16.5 (**fig. S5G**). Similarly, inducible deletion of *Gpx4* during the postnatal cornea lymphatic vascular developmental stage i.e. from p0 to p4, reduced lymphatic growth at p10 (**Fig. 3G**), while deletion of *Gpx4* during the regression phase (p16-p18) (**fig. S5H**) had no impact in lymphatic density at p22 as lymphatic endothelial cells naturally regress through ferroptosis (See **Fig. 1**).

**Fig. 3.**
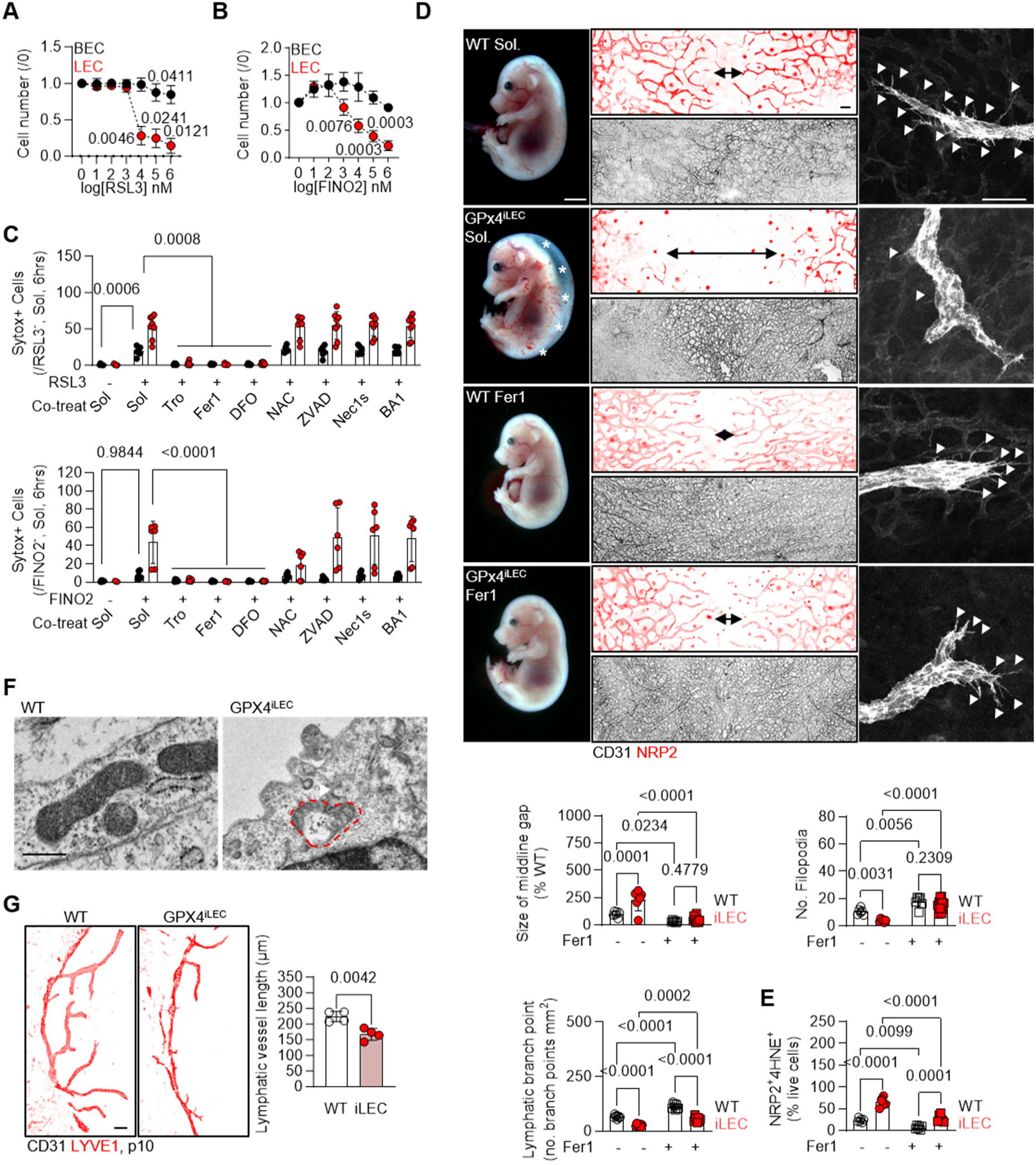
Lymphatic vessel growth and survival depend on GPX4. (**A-B**) Relative cell number of human lymphatic (LEC) and blood (BEC) endothelial cells in response to increasing concentrations of the GPX4 inhibitor RSL3 (A) or FINO2 (B). n=3/ group in 3 independent experiments. Kolmogorov-Smirnov normality test. One way Anova, Tukey’s post hoc analysis. (**C**) Relative Sytox positive cells as in panel (A), cultured in the presence or absence of RSL3 (upper panel, 5 μmol/L), or FINO2 (lower panel, 50 μmol/L) and simultaneously exposed to Trolox (Tro, 100 μmol/L), Ferrostatin1 (Fer1, 10 μmol/L), Deforoxamine (DFO, 100 μmol/L), N-acetyl-L-cysteine (NAC, 1 mmol/L), Z-VAD-FMK Caspase inhibitor (ZVAD, 50 μmol/L), Necrostatin (Necs1, 10 μmol/L) or Bafilomycin 1 (BA1, 1 nmol/L) for 6 hours respectively. n=3/ group in 3 independent experiments. Two way Anova, Dunnet’s multiple comparisons test. (**D**) Stereomicrographic images of E15.5 embryos and immunofluorescent images of CD31 (black) blood and NRP2 (red) lymphatic dermal endothelial cells in PROX1 Cre^-^ (WT) or Cre^+^ (GPx4^iLEC^) mice treated with tamoxifen from E9.5-E12.5 to induce genetic deletion of lymphatic endothelial cell *Gpx4*. In some cases, pregnant mice received simultaneously Ferrostatin 1 from E10.5 up to E14.5 (Fer1, 2 mg/kg/day, gavage). Right panel shows NRP2 (gray) lymphatic filopodia in high magnification. Relative quantification of middline gap as % to WT, number of filopodia and lymphatic branch points in the same embryos. Asterisks: murine edema. Black arrows: indicator of middline gap. White arrows: filopodia. Scale bars = 1 mm, 100 μm and 50 μm. n=6-11 embryos/ group from 5 independent births. Two way Anova, Fischer’s multiple comparison test. (**E**) Percentage of NRP2^+^ 4HNE^+^ live cells in samples as in panel (D). n=6-8 embryos/ group. Two way Anova, Fischer’s multiple comparison test. (**F**) Transmission electron microscopy of the dermal lymphatic endothelial cells from embryo skin at E15.5 in samples as in panel D. Arrows: mitochondrial morphology; dashed red lines: mitochondria. Scale bars = 500 nm. (**G**) Immunostaining of LYVE1 (red) in postnatal day (p) 10 murine cornea after inducing lymphatic specific deletion of GPX4 (iLEC) from p0 to p6, and respective quantification of lymphatic vessel length. n=4 corneas/ group. Scale bars = 100 μm. Kolmogorov-Smirnov normality test, Unpaired Student’s *t* test.

### GPX4 degradation is responsible for lymphatic endothelial cell ferroptosis sensitivity

How do lymphatic but not blood endothelial cells exhibit GPX4-dependent ferroptosis sensitivity? Initially, we examined the possibility that the two cell types show distinct redox gene signatures despite originating from the same precursors. RNA sequencing analysis and comparison between dermal blood and lymphatic endothelial cells, revealed that blood endothelial cells in general hold a more anti-oxidant transcriptional signature (**fig. S6A**); with increased transcriptional levels of the ferroptosis protecting and glutathione metabolism genes (**fig. S6B**); yet, such transcriptional signatures did not suffice to explain the strong GPX4 dependent sensitivity to ferroptotic cell death seen in the lymphatic vasculature.

Tracing 13C labelled cystine in dermal blood and lymphatic endothelial cells, revealed that lymphatics contained a low GSH/GSSG ratio, which is indicative of decreased GPX4 function (**Fig. 4A**). Indeed, GPX4 protein levels were lower in the human dermal lymphatic than in blood endothelial cells (**Fig. 4B**). Inhibition of protein synthesis resulted in a time dependent decrease of GPX4 levels only in the lymphatic but not blood endothelial cells (**Fig. 4C**), which was reversed by the addition of the proteasomal degradation inhibitor MG132 but not the autophagy inhibitor Bafilomycin A (**Fig. 4C**). These observations led to the hypothesis that protein stability mechanisms linked to the ubiquitin-proteasome-system (UPS) are potentially responsible for the regulation of GPX4 expression. Therefore, we reasoned that differential ubiquitin dependent degradation mechanisms were present in the two endothelial cell types. Fitting with this, lymphatic endothelial cells showed increased K48 GPX4 polyubiquitination compared to the blood endothelial cells (**fig. S6C**); which is indicative of an active GPX4 proteoasomal degradation mechanism (*53*).

**Fig. 4.**
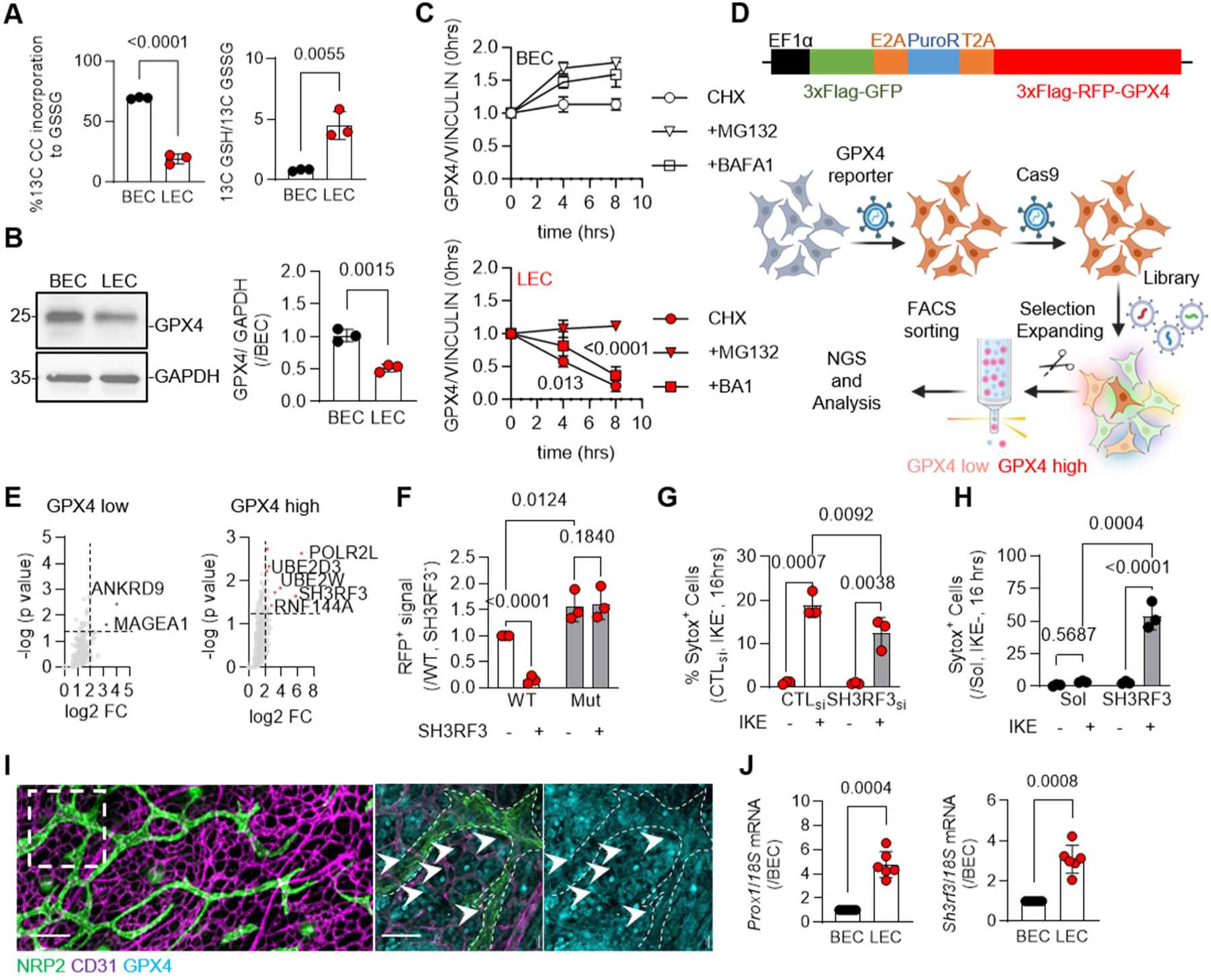
GPX4 degradation surveils ferroptotic mechanisms in the lymphatic endothelium. (**A**) Percentage of 13C Cystine (CC) incorporation to GSSG and 13C GSH/13C GSSG ratio in human blood (BEC) and lymphatic (LEC) endothelial cells. n=3/group. Shapiro-Wilk normality test, unpaired Student‘s t test. (**B**) Representative immunoblot of GPX4 and GAPDH and respective quantification in samples as in panel (A). n=3/group. Shapiro-Wilk normality test, unpaired Student‘s t test. (**C**) Response of GPX4 to cycloheximide (CHX, 10 μg/mL) treatment for up to 8 hours in the presence or absence of the proteasomal inhibitor MG132 (10 μmol/L) and the autophagy inhibitor Bafilomycin A (BA1, 320 nmol/L) in human blood (BEC) and lymphatic (LEC) endothelial cells. n=4/group. Two way Anova, Dunnet’s multiple comparisons test. (**D**) Schematic showing the GPX4 protein stability reporter construct and the CRISPR Cas9 strategy for the identification of human ubiquitin ligases and deubiquitinases responsible for its regulation. (**E**) Volcano plots showing the respective gRNA-gene targets regulating GPX4 stability; negatively selected following the CRISPR screening presented in panel (D). ANKRD9:Ankyrin repeat domain-containing protein 9; MAGEA1: Melanoma-associated antigen 1; POLR2L: DNA-directed RNA polymerases I, II, and III subunit RPABC5; UBE2D3: Ubiquitin-conjugating enzyme E2 D3; UBE2W: Ubiquitin-conjugating enzyme E2 W; SH3RF3: E3 ubiquitin-protein ligase SH3RF3; RNF144A; Ring Finger Protein 144A. (**F**) Relative RFP positive signal (GPX4 high cells) following in wild type or K107, K162 and K167 to R triple GPX4 reporter mutants in the presence or absence of 1 µg purified SH3RF3 protein. n=3/group (in 3 technical replicates). One way Anova, Tukey’s multiple comparisons test. (**G**) Relative Sytox positive human lymphatic endothelial cells treated with a control siRNA (CTL_si_) or a siRNA directed against SH3RF3 (SH3RF3_si_) in the presence or absence of IKE (10 μmol/L) for 16 hours. n=3/group (in 3 technical replicates). Two way Anova, Fischer’s multiple comparison test. (**H**) Relative Sytox positive human blood endothelial cells treated with purified SH3RF3 protein in the presence or absence of IKE (10 μmol/L) for 16 hours. n=3/group (in 3 technical replicates). Two way Anova, Fischer’s multiple comparison test. (**I**) Immunostaining of GPX4 (cyan) in NRP2 (green) lymphatic and CD31 (magenta) blood endothelial cells on E15.5 wild type embryo skin. Scale bars = 150 μm and 20 μm. (**J**) Relative *Prox1* and *Sh3rf3* mRNA levels in murine dermal blood (BEC) and lymphatic (LEC) endothelial cells isolated from E15.5 wild type embryo skin. n=6/group. Kolmogorov-Smirnov normality test, paired Student‘s t test.

To identify potential ligases and deubiquitinases responsible for determining the stability of GPX4, a fluorescent reporter construct (*54*) that allows live cell monitoring of intracellular GPX4 protein stability was generated and used in a gRNA CRISPR screening utilizing a newly synthetized deubiquitinase and ligase library (*55*) (**Fig. 4D**). This approach revealed that GPX4 expression was sensitive to the depletion of the E2 ubiquitin conjugating enzymes UBE2D3 and UBE2W, as well as the E3 ubiquitin ligase SH3RF3 (**Fig. 4E, Table S1**). Although both blood and lymphatic endothelial cells differentially expressed many deubiquitinases and ubiquitin ligases, all three USP related hit proteins were highly expressed in the lymphatic but not blood human endothelium (**fig. S6D**). Given that SH3RF3 is a RING finger like E3 ubiquitin ligase, implicated in numerous cellular processes (*56*), we next focused on the impact of SH3RF3 on GPX4 protein stability. Several lysines (K) of GPX4 have been previously reported as potential ubiquitination sites (*57*) and were subsequently mutated to Arginine (R) K107, K162 and K167 Lysine (K) residues to Arginine (R). Wild type GPX4, but not the GPX4 3R mutant, was destabilized upon addition of active SH3RF3, thereby validating GPX4 as a substrate of this E3 ligase and a UPS target (**Fig. 4F**). Functionally, silencing of SH3RF3 in the lymphatic endothelium reduced ferroptosis sensitivity to IKE (**Fig. 4G**), while addition of purified SH3RF3 to blood endothelial cells increased IKE mediated cell death (**Fig. 4H**). In alignment with our *in vitro* findings, *in vivo* analysis of developing lymphatic vessels in the murine skin at E15.5 revealed reduced GPX4 expression compared to neighboring blood endothelium (**Fig. 4I**), and high SH3RF3 levels (**Fig. 4J**). Taken together, we identified a ubiquitin-proteasome-system related molecular mechanism that differentially regulates GPX4 protein levels in lymphatic vessels, ultimately allowing selective regression of the lymphatic endothelium while keeping the neighboring blood endothelium intact.

### Membrane PUFA enrichment promotes ferroptosis sensitivity in the lymphatic endothelium

Susceptibility to ferroptosis due to loss of the ferroptosis suppressive pathways involving cystine and glutathione utilization might suffice to explain partially the lymphatic endothelial cell death responses, however, when such ferroptotic suppressing pathways are tampered, cells render themselves sensitive to ferroptosis only in the presence of relevant lipid peroxidation acceptors. Among those membrane phospholipids and in particular those with polyunsaturated fatty acid (PUFA) tails are considered good acceptors of reactive oxygen species (*58, 59*). In addition, peroxisomal generated PUFA ether phospholipids (ePLs) are significant substrates for lipid peroxidation, resulting in the induction of ferroptosis (*60*). In contrast to PUFAs, saturated and monounsaturated fatty acids (SFAs, MUFAs) are resistant to peroxidation and excess MUFAs can even protect cells from ferroptosis (*61, 62*). Lipidomic analysis revealed that lymphatic endothelial cells were enriched in global PUFA glycerolphospholipids (**Fig. 5A**, **Table S2**). Approximately 56% less phospholipids with one SFA tail, 50% less phospholipids with two SFA tails, 20% more phospholipids with one PUFA tail and 50% more phospholipids with two PUFA tails were detected in dermal lymphatic endothelial cells compared to dermal blood endothelial cells (**Fig. 5B**). In particular, lymphatic endothelial cells showed reduced abundance of glycerolphosphatidylcholine and ethanolamine with two SFA tails as well as significant less incorporation of SFA and MUFA tails (**Fig. 5C, Table S3**). In contrast, increased presentation of glycerolphospholipids with one or two PUFA tails in the lymphatic endothelium was observed (**Fig. 5C, Table S3**). In total, the PUFA to SFA ratio in both phosphatidylcholine (**Fig. 5D**) and phosphatidylethanolamine (**Fig. 5E**) were increased approximately 3-4 fold in lymphatic vs. blood endothelial cells. Notably, phospholipids with two PUFA tails exceeded 15-35% of total phosphatidylcholine and phosphatidylethanolamine in the lymphatic but were significantly less in the blood endothelium (**Fig. 5F, 5G**). Ether linked glycerolipids (sn1 position) originating from peroxisomes (*60*) were also enriched in the lymphatic endothelial cells, with particularly ether phosphatidylcholine (ePC) being significantly abundant in the lymphatic compartment and decorated with PUFA tails (**Fig. 5H**, **Table S4**). Such heterogeneity could be attributed to the differential expression of the ether phospholipid generating mRNAs (**Fig. 5I**). Notably cholesterol esterified fatty acids were found enriched in the blood endothelium (**fig. S7A**), in line with the increased expression of the acetyl-CoA acetyltransferase 1 (ACAT1) enzyme (**fig. S7B**), an accumulation that potentially neutralizes their capacity to react with radicals (*60*). Diacyl glycerolipids (DAG) also showed differences between the studied groups, with lymphatic endothelial cells enriching primarily for PUFA2 DAGs as well as SFA/MUFA DAGs; while triacyl glycerolipids (TAG) where reduced in the lymphatic endothelium (**fig. S7C**).

**Fig. 5.**
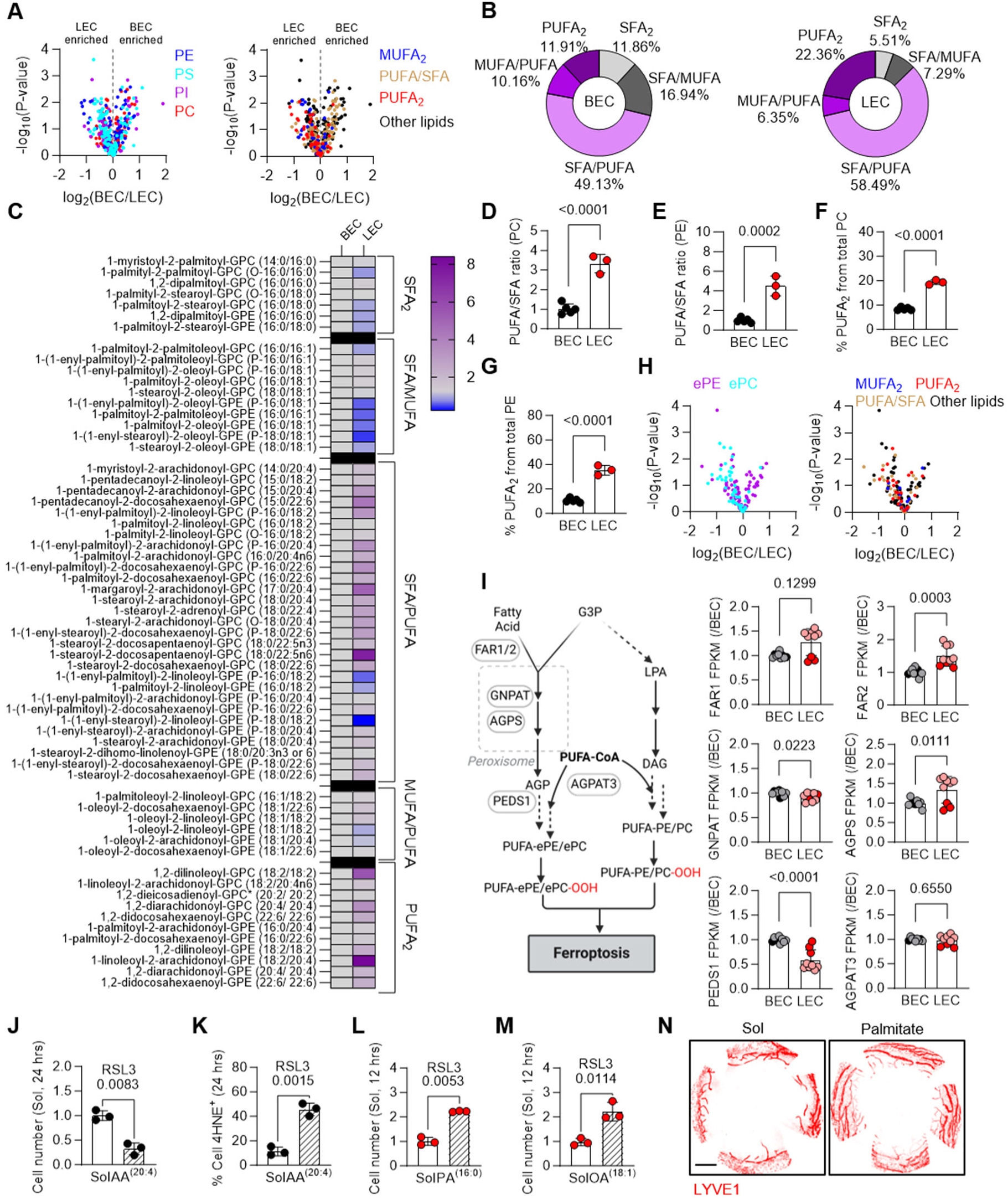
Lymphatic endothelial cell sensitivity to ferroptosis is driven by increased PUFA2 phospholipid incorporation. (**A**) Volcano plots showing glycerolphospholipid clusters (left panel) and their respective fatty acid tail clusters (right panels) detected in human dermal blood (BEC) and lymphatic (LEC) endothelial cells. n=3/ group. Unpaired Student’s t test. (**B**) Percentage of fatty acid glycerolphospholipid tails from total glycerolphospholipids detected in human blood (BEC) and lymphatic (LEC) endothelial cells. (**C**) Heatmap showing glycerolphosphatidylcholine (GPC) and glycerophosphatidylethanolamine (GPE) phospholipids detected in human blood (BEC) and lymphatic (LEC) endothelial cells. n= 3-5/ group. (**D**) PUFA/SFA ratio of phosphatidylcholine (PC) in samples as in panel (C). n= 3/ group. Shapiro-Wilk normality test, unpaired Student’s t test. (**E**) PUFA/SFA ratio of phosphatidylethanolamine (PE) in samples as in panel (C). n= 3/ group. Shapiro-Wilk normality test, unpaired Student’s t test. (**F**) Percentage of glycerophospholipids with two PUFA tails (PUFA_2_) from total PC in samples as in panel (C). n= 3/ group. Shapiro-Wilk normality test, unpaired Student’s t test. (**G**) Percentage of glycerophospholipids with two PUFA tails (PUFA_2_) from total PE in samples as in panel (C). n= 3/ group. Shapiro-Wilk normality test, unpaired Student’s t test. (**H**) Volcano plots showing ether phospholipids clusters (left panel) and their respective fatty acid tail clusters (right panels) detected in human dermal blood (BEC) and lymphatic (LEC) endothelial cells. n=3/ group. Unpaired Student’s t test. (**I**) Schematic showing ether lipid generating enzymes and relative transcriptional levels of fragments per kilobase million in human blood (BEC) and lymphatic (LEC) endothelial cells. n=9/group. Mann-Whitney test, Kolmogorov-Smirnov and D’Agostino & Pearson normality tests, unpaired Student‘s t test. Light colors represent juvenile while dark colors adult cells. FAR1/2: Fatty Acyl-CoA Reductase 1/2; GNPAT: Glyceronephosphate O-Acyltransferase; AGPS: Alkylglycerone Phosphate Synthase; PEDS1: Plasmanylethanolamine Desaturase 1; AGPAT3: 1-Acylglycerol-3-Phosphate O-Acyltransferase 3; G3P: glycerol-3-phosphate; 1-O-Alky-glycerol-3-phosphate; LPA: Lysophosphatidic acid; DAG:Diacylglycerol. (**J**) Relative cell number of human blood endothelial cells following 24 hour treatment with Solvent (Sol) or the PUFA Arachidonic Acid (AA, 100 µmol/L) and subsequently exposed to RSL3 (5 µmol/L, 24 hrs). n=3/group. Shapiro-Wilk normality test, unpaired Student’s t test. (**K**) Percentage of 4HNE^+^ cells in samples as in panel J. n=3/group. Shapiro-Wilk normality test, unpaired Student’s t test. (**L**) Relative cell number of human lymphatic endothelial cells following 24 hours treatment with Solvelnt (Sol) or the SFA Palmitic Acid (PA, 100 µmol/L) and subsequently exposed to RSL3 (5 µmol/L, 24 hrs). n=3/group. Shapiro-Wilk normality test, unpaired Student’s t test. (**M**) Relative cell number of human lymphatic endothelial cells following 24 hours treatment with Solvent (Sol) or the MUFA Oleic Acid (OA, 100 µmol/L) and subsequently exposed to RSL3 (5 µmol/L, 24 hrs). n=3/group. Shapiro-Wilk normality test, unpaired Student’s t test. (**N**) Immunostaining of LYVE1 (red) lymphatic endothelial cells on postnatal day 16 corneas after receiving daily treatment with solvent (Sol) or the SFA Palmitic acid (PA, 5 μL of 500 μmol/L) since postnatal day 14. Scale bar= 500 µm.

To investigate potential regulators of such differential phospholipid signature in the two cell types, we next evaluated the transcriptional expression of the major fatty acid regulating genes (**fig. S8A**). Interestingly in line with the reduced SFA availability, dermal lymphatic endothelial cells showed reduced transcription of the SFA elongation (ELOVL fatty acid elongase 6), synthesis (fatty acid synthase) and desaturation (stearoyl CoA desaturase) compared to dermal blood endothelial cells (**fig. S8B**). In contrast, increased fatty acid binding protein 4 (FABP4) and the fatty acid transporting protein CD36 transcripts were detected in the dermal lymphatic endothelial cells (**fig. S8C**), which have been implied to serve as albumin bound PUFA importing proteins (*63*). Indeed, CD36 expression was functionally relevant to the sensitivity of the lymphatic endothelial cells to ferroptosis as inhibition of CD36 using sulfosuccinimidyl oleate rendered lymphatic endothelial cells resistant to RSL3 mediated ferroptosis (**fig. S8D**). On the other hand, overexpression of CD36 in blood endothelial cells rendered them sensitive to RSL3-induced ferroptotic death (**fig. S8E**).

Although, no major differences were observed in the transcriptional levels of the EVOLV1 and 5 elongases (**fig. S9A**) as well as phospholipases (**fig. S9B**), a significant increase in the fatty acid desaturases (FADS) 2 and 3 was observed in the dermal lymphatic endothelial cells (**fig. S9C**). Members of the acyl-CoA synthetase long chain family (ACLS) potentially shape the cellular lipid composition that dictates sesnsitivity to ferroptosis (*64*). Of the ACLS genes only the expression of ACLS4 was significantly different between blood and lymphatic endothelial cells (**fig. S9D**). Lentiviral deletion of ACLS3, 4 and 5 but not ACLS1 prolonged cell survival in response to GPX4 pharmacological inhibition in the lymphatic endothelial cells, without affecting the death of the blood endothelial cells (**fig. S9E**); while overexpression of ACLS5 in the blood endothelium increased susceptibility to RSL3 mediated ferroptotic cell death (**fig. S9F**). In addition, significant reduction of the majority of the mRNAs from proteins responsible for the PUFA incorporation was observed (**fig. S9G**). Taken together, these data indicate a simultaneous contribution of PUFA activation and incorporation mechanisms that increase the lymphatic endothelial cell membranes sensitivity to lipid peroxidation. However, transcriptional differences in the acyl-CoA synthetases and PUFA incorporation mRNAs, was not the only difference explaining the reduced PUFA incorporation in the blood endothelial cells making them resistant to ferroptosis. PUFAs and particular arachidonic acid serve as precursors of multiple signalling cascades to generate leukotriens, bioactive eicosanoids as well as prostanoids (*65*) (**fig. S9H**). Blood endothelial cells showed also increased metabolic flux of the arachidonic acid to the prostaglandins E2 and Fa2 (**fig. S9I**) following increased expression of the two rate limiting enzymes PTGS1 and 2 (**fig. S9J**).

To confirm that the increased availability of lipid peroxidation accepting fatty acids, was the ultimate limiting factor of ferroptosis resistance in blood endothelial cells alongside the tampered cystine/glutathione availability, we manipulated the fatty acid consumption in proliferating blood and lymphatic endothelial cells. Phenotypically, addition of a fatty acid free BSA conjugated PUFA i.e. arachidonic acid in proliferating blood endothelial cells, increased RSL3 mediated death (**Fig. 5J**) and lipid peroxidation (**Fig. 5K**). In contrast, addition of a fatty acid free BSA conjugated SFA i.e. palmitic acid (**Fig. 5L**) or MUFA i.e. oleic acid (**Fig. 5M**) in proliferating lymphatic endothelial cells, inhibited RSL3 mediated ferroptotic cell death *in vitro*. These observations are in line with previous reports that oleic acid availability within the lymph fluid protects melanoma cells from ferroptosis (*66*). *In vivo* supplementation of a fatty acid free BSA conjugated palmitic acid in the postnatal cornea during the initiation of the regression phase i.e. from p14-16 increased lymphatic growth and halted regression (**Fig. 5N**). Taken together, these data indicate that loss of cystine/glutathione dependent ferroptotic suppresion mechanisms and differential membrane PUFA enriched lipid composition, work synergistically to fine-tune lymphatic growth and regression.

### Induction of ferroptosis after lymphatic injury, halts lymphatic hyperplasia and shapes immune cell responses

Lymphangiogenesis and angiogenesis are both activated in conditions following injury of the cornea and contribute to corneal transplant failure (*67, 68*). Therefore, next, we evaluated whether manipulating lymphatic endothelial cell ferroptosis would be a potential target to regulate lymphatic growth after cornea injury. Inducible genetic deletion of GPX4 in the lymphatic vasculature resulted in reduced lymphatic growth 7 days after cornea injury (**Fig. 6A**). This was accompanied by reduced lymphatic vessel area (**Fig. 6B**), branching points (**Fig. 6C**), length (**Fig. 6D**) as well as blood vessel growth (**Fig. 6E**). All of these effects were rescued by the addition of the ferroptosis inhibitor, ferrostatin 1. Similar results were obtained, when wild type mice were treated with SLC7A11 or GPX4 inhibitors (**fig. S10**), with an overall reduction of lymphangiogenic responses observed in the injured cornea, rescued by the addition of ferrostatin 1. Notably, lymphatic induction of ferroptosis resulted in reduced immune cell infiltration on day 3 post injury, with a particularly strong impact on neutrophils, CD4^+^ T cells and monocyte infiltration (**Fig. 6F**). These data suggest a role for physiological lymphatic ferroptosis in altered immune response after injury, characterized by a dominant impact on minimizing monocyte infiltration that may indicate non-phlogistic tissue remodelling. In contrast, reduced susceptibility to ferroptosis achieved by the addition of the SFA palmitic acid for 7 constitutive days, protected against lymphatic loss and potentiated lymphangiogenesis in the injured corneas (**Fig. 6G**). Addition of the PUFA arachidonic acid, halted lymphagiogenic responses and lymphatic overgrowth following cornea injury (**Fig. 6G**).

**Fig. 6.**
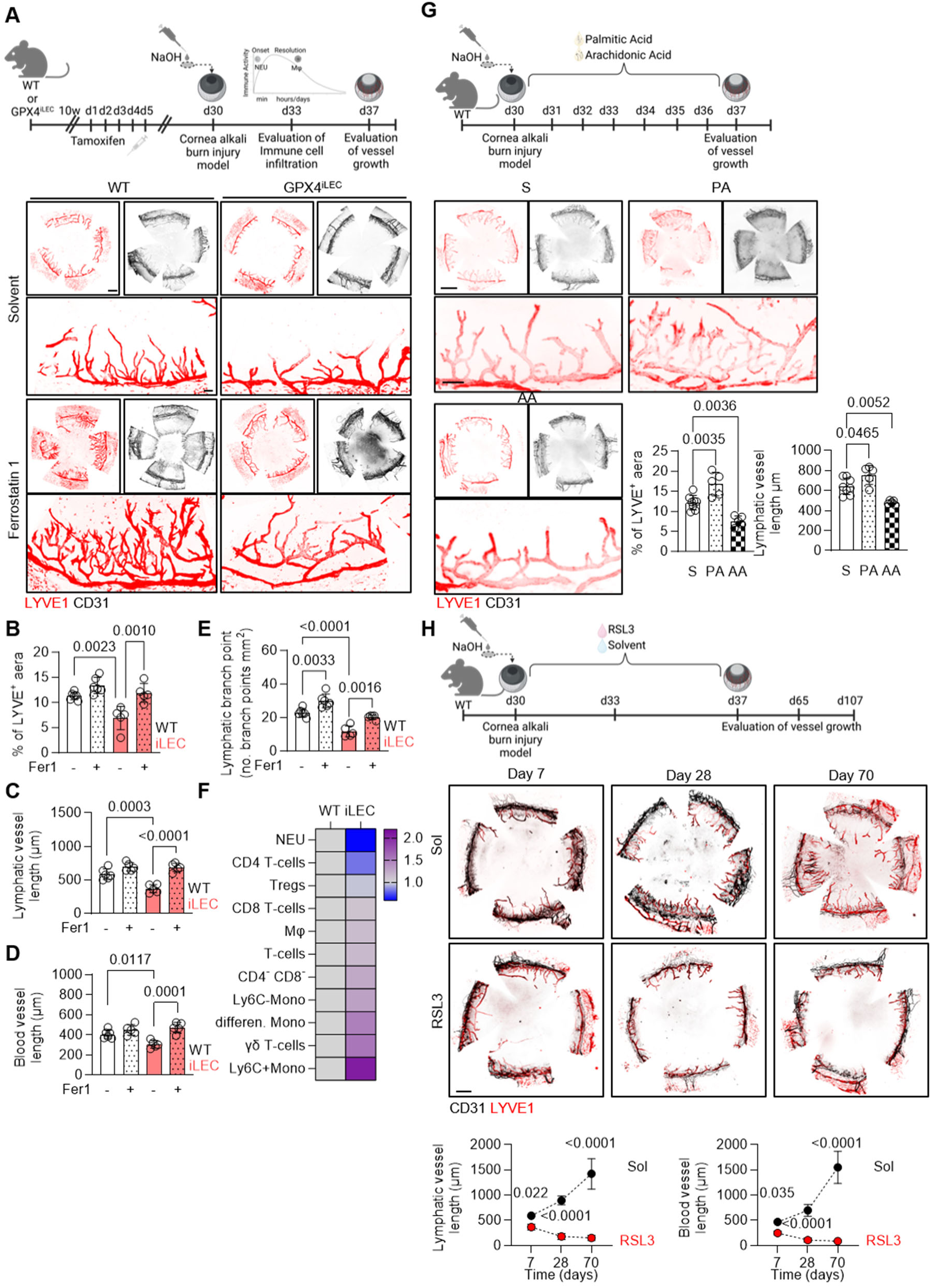
Manipulating ferroptosis impacts on lymphatic vessel growth and immune cell infiltration after injury. (**A**) Representative immunostaining of LYVE1 (red) lymphatic and CD31 (black) blood endothelial cells in adult corneas from wild type (WT) or lymphatic specific inducible *Gpx4* deleted mice (iLEC) following alkali burn cornea injury for 7 days. Scale bars = 500 µm and 100 µm. Corneas were treated with solvent (Sol) or Ferrostatin 1 (Fer1, 50 μmol/L) starting at the day of injury and daily up to 7 days post-injury. (**B**) Percentage of LYVE1 positive area in samples as in panel (A). n= 5-6/ group. Kolmogorov-Smirnov normality test. One way Anova (Tukey’s multiple comparison test). (**C-D**) Lymphatic (C) and blood (D) vessel length in µm samples as in panel (A). n= 5-6/ group. Kolmogorov-Smirnov normality test. One way Anova (Tukey’s multiple comparison test). (**E**) Lymphatic number of branch points/mm^2^ in samples as in panel (A). n= 5-6/ group. Kolmogorov-Smirnov normality test. One way Anova (Tukey’s multiple comparison test). (**F**) Immune cell infiltration in the cornea at day 3 post-injury in samples as in panel A following flow cytometry analysis. n= pool of 8 corneas/ group. (**G**) Immunostaining of LYVE1 (red) lymphatic and CD31 (black) blood endothelial cells in adult corneas from wild type mice treated with solvent (S), arachidonic acid (AA, 5 µl, 500 µmol/L) or palmitic acid (PA, 5 µl, 500 µmol/L) following alkali burn cornea injury for 7 days and respective quantification of the LYVE1 percentage and lymphatic vessel length. n=5-10/ group. Scale bar = 500 µm and 100 µm. Kolmogorov-Smirnov normality test. One Way Anova, Tukey‘s multiple comparison test. (**H**) Representative immunostaining of lymphatic (LYVE1) and blood (CD31) endothelial cells in adult corneas from wild type mice treated with RSL3 (10 μmol/L) or Solvent (Sol.) between 0 and 7 days post-injury and monitored for up to 70 days and respective lymphatic and blood vessel length quantification. n=6/ group. bar = 500 µm. Kolmogorov-Smirnov test. Paired Student‘s t test for each time point.

In an attempt to prevent excessive lymphangiogenesis after injury, we administrated three doses of the GPX4 inhibitor, RSL3, at day 0, day 3 and day 7 post injury (**Fig. 6H**). Interestingly, ferroptosis induction at those specific time points resulted in fully normal corneal lymphatics already at day 14 after injury, while solvent treated corneas exhibited excessive regeneration for up to 70 days after injury (**Fig. 6H**).

## Discussion

Lymphatic dysfunction is increasingly recognized as a key contributor to a broad spectrum of diseases, including lymphedema, neurological and neurodegenerative disorders, inflammatory bowel disease, cardiovascular disease, metabolic syndrome, and glaucoma. Conversely, pathological lymphangiogenesis is associated with inflammatory conditions, cancer progression, and transplant rejection (*15*). For instance, successful transplantation of lymphatic enriched organs, like the cornea, heavily relies on adjusting lymphatic growth (*67, 68*). Despite extensive studies on lymphangiogenic pathways, from embryonic lymphatic specification to adult vessel remodeling, therapeutic strategies targeting lymphatic regression or endothelial cell death remain unexplored. Only limited knowledge on lymphatic regression in genetic models overexpressing a soluble VEGFR3 form has been reported, which suggest that inhibition of VEGF-C/VEGF-D signalling induces a regression of already formed lymphatic but not blood vessels (*69*). Here we show that ferroptosis is a physiological mechanism regulating lymphatic endothelial cell turnover during developmental processes and after injury. Our data reveal that ferroptosis regulates lymphatic endothelial cell growth and regression through intrinsic cell-type specific pathways that affect both ferroptosis suppressive pathways as well as phospholipid membrane composition. Interfering with such mechanisms in the lymphatic endothelium is proven beneficial in preclinical mouse models of lymphatic injury, without compromising the blood vasculature.

Although ferroptosis is considered one of the most ancient form of death mechanisms, still its natural inhibitors or inducers remain elusive. Ferroptosis is ultimately driven by the peroxidation of specific membrane lipids and ferroptosis-resistance is tightly linked to membrane incorporation of polyunsaturated fatty acids (*64, 70–72*) and structure of specific phospholipid tails (*58, 73*). Sensitivity to ferroptosis is also regulated by glutathione biosynthesis (*74*). Increased efflux of glutathione (*75*), cysteine deoxygenase 1 mediated deletion of cysteine (*76*), or chemical induced GPX4 degradation (*44*), all increase sensitivity to ferroptosis. Notably, chaperone mediated autophagy and GPX4 degradation, has also been proposed to regulate ferroptosis sensitivity (*77*). Although, the link between protein ubiquitination and ferroptosis sensitivity has only started to be investigated (*78*), and the impact of GPX4 post translational modifications in the regulation of GPX4 protein stability is minimally studied, a broad-spectrum deubiquitinase inhibitor, has been just reported to increase GPX4 protein degradation in non-small cell lung cancer cells (*79*). Here, we report a previously uncharacterized GPX4 degradation mechanism governed by a cell-type specific Ring finger E3 ligase that responds to the maturation and growth status of lymphatic endothelial cells. This endogenous pathway triggers ferroptosis without exogenous inducers, establishing a new regulatory axis, which relies on the capacity of the cell to degrade the rate limiting enzyme GPX4, for maintaining the lymphatic homeostasis.

Unexpectedly, we discovered that lymphatic endothelial cells differ from blood endothelial cells, although rising mostly from the same progenitors, in terms of PUFA availability and incorporation to phospholipids. Lymphatic endothelial cells are uniquely enriched in phospholipids containing two PUFA chains (PL-PUFA2_s_), a rare lipid species linked to heightened ferroptosis sensitivity (*58*). Lymphatic dermal endothelial cells were found to significantly enrich PL-PUFA_2_ species in their membranes to a great extent compared to other mammalian endothelial cells. This is attributed to the increased import and activation of arachidonic acid PUFAs in the lymphatic endothelium, coupled with reduced utilization of the arachidonic acid towards the prostaglandin biosynthesis. Interestingly, fatty acids, glutathione and free iron content in lymph and blood fluids have been shown to be different resulting in extrinsic regulation of ferroptosis to cancer cells (*66*). Although, such differences could also impact directly the ferroptotic responses of the blood and lymphatic endothelium *in vivo*, here we highlight additional specific fatty acid mechanisms that impact on membrane phospholipid composition to alter cellular decisions against ferroptosis sensitivity and resistance. Notably, increased accumulation of phospholipids with two PUFA tails enhanced the ferroptotic sensitivity of the lymphatic endothelium, and addition of SFAs in the developing or injured lymphatics reinstated ferroptosis resistance in the specific vascular bed. Thus, potential manipulation of fatty acid availability during lymphatic injury, could serve as a strategy to enhance lymphagiogenic responses or promote lymphatic regression.

Our findings also have implications in transplantation biology, where allograft rejection remains a leading cause of graft failure beyond the first year, despite immunosuppression. Impaired lymphatic regeneration in all solid organ transplants, due to surgical disconnection and technical limitations, contributes to poor graft outcomes (*80*). Emerging studies indicate that modulating lymphangiogenesis may mitigate transplant arteriosclerosis (*81*) and improve corneal transplant success by preserving tissue transparency (*82*). As the organ-specific roles of lymphatics become clearer, a reconceptualization of their function in solid organ transplantation is warranted to assess whether therapeutic modulation can enhance clinical outcomes. In the present study, we identified that by promoting or inhibiting ferroptosis, we could manipulate the growth of the lymphatic vasculature offering a tool for modulating vascular remodeling in developmental or transplantation settings. Interestingly, we also observed that lymphatic endothelial cell ferroptosis, influences local immune responses to injury, a finding that contributes to understanding how physiological cell death mechanisms regulate immunosurveillance of specific organs. Potential translation of the present findings to the cornea transplantation procedures would benefit allograft acceptance. In addition, based on the current evidence presented here, concerns should be raised on the studies utilizing ferroptosis inducers as anti-cancer strategies, given the potential direct events to the lymphatic vascular bed survival and function.

Altogether, our study reveals a physiological role of ferroptosis in lymphatic vascular remodeling, driven by intrinsic metabolic and ubiquitin-regulated pathways, and uncovers new mechanisms to target lymphatic growth and homeostasis. Given the broad implications of lymphatic dysfunction across multiple pathologies, these insights open promising opportunities for therapeutically modulating lymphatic growth in both regenerative medicine and disease contexts.

## Data Availability

All data associated with this study are present in the paper or Supplementary Materials. Requests for material should be sent to the corresponding author.

## Acknowledgements

The authors are indebted to Britta Heckmann for expert technical assistance. We thank the Core Facilities Pre-Clinical Models, LIMA (Live Cell Imaging) and FlowCore for their assistance. All graphics were designed with Biorender.

## Funding

This work was supported by the Deutsche Forschungsgemeinschaft (CRC1531/1 Project A02 to S.-I.B., Project B03 to J.H and I.F., Project ID 456687919; CRC1366/2 Project B01 to S.-I.B., Project A03 to G.D., Project ID 39404578; Project F08 to S.-I.B., Project F06 to A.S, Project F03 to M.K, Project ID 259130777; the Emmy Noether Programme BI 2163/1-1 to S.-I.B.; the Hector Fellow Academy to S.-I.B. and the BI 2482/3-1 to S.-I.B.); the National Natural Science Foundation of China, Project IDs; 82070501, 82271479 and 32350021 to J.H.; M.-K.D. was supported with a scholarship from Onassis Foundation.

## Author Contributions

Conceptualization: S.-I.B., J.H. and M.-K.D. Study design: S.-I.B., J.H, M.-K.D and D.W. Supervision: S.-I.B and J.H. Funding acquisition: S.-I.B. and J.H. Methodology: M.-K.D., D.W., Y. C., Y.D., J.W., R.X., D.S., B.A., J.M., L.G., C.X., R.C., Y.M., M.K., A.W. Sequencing and Bioinformatics: S.G. Writing original draft: S.-I.B. Conceptual advice: I.F., M.K., G.D., A.W., T.P.D., A.S., I.D., A.L. Writing-review & editing: All authors.

## Competing interests

Authors declare that they have no competing interests.

## Supplementary Figure Legends

**Fig. S1.**
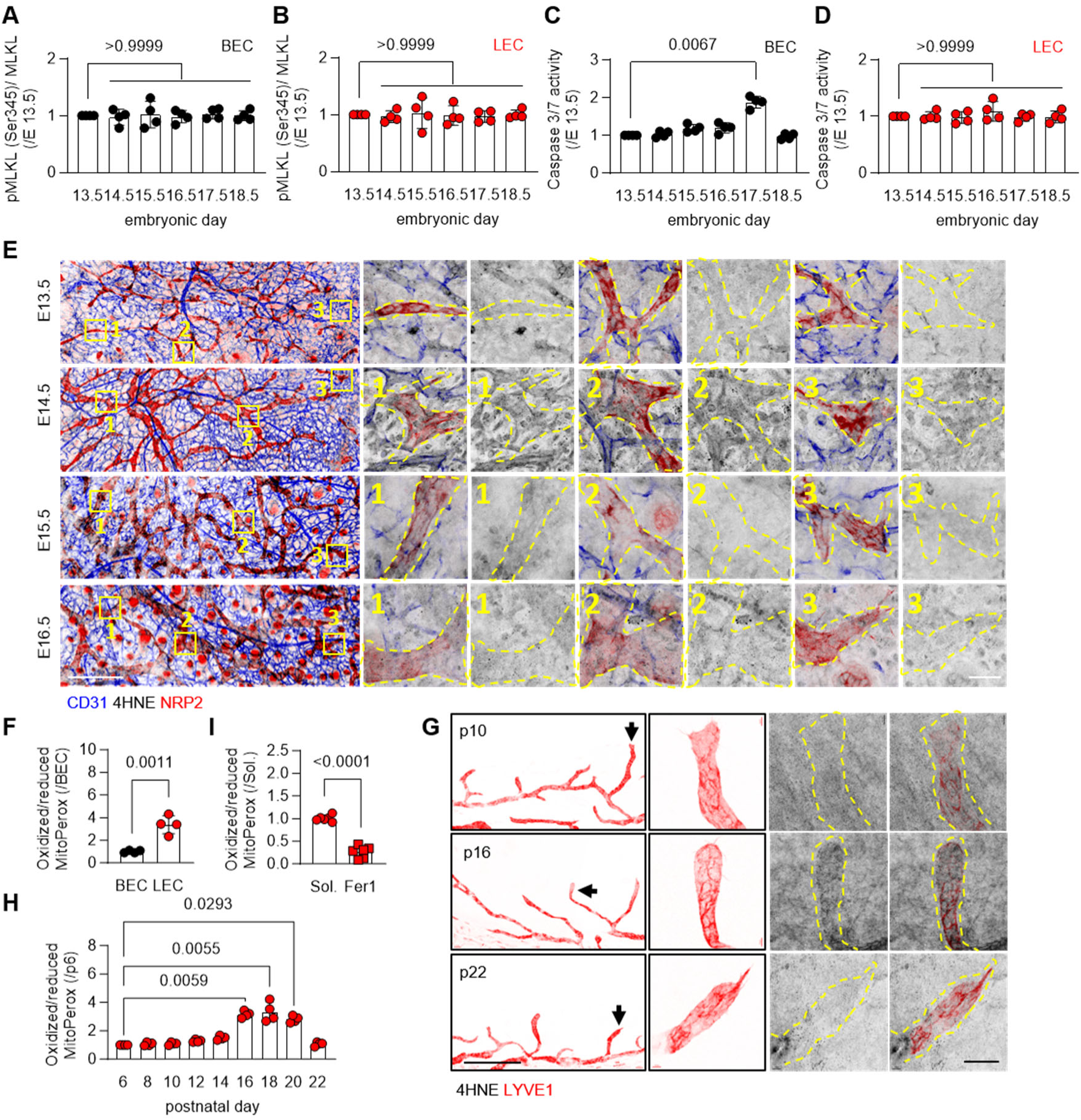
Lipid peroxidation marks growing lymphatic but not blood endothelial cells. (**A-B**) Respective quantification of phosphorylated MLKL (Ser345) to total MLKL ratio in PROX1^-^CD31^+^ blood (BEC) and PROX1^+^ lymphatic (LEC) dermal endothelial cells isolated from samples as in Figure 1, panel (A). n=4/ group, with each sample a pool of 6-10 embryos, Kruskal-Wallis (Dunn’s multiple comparison test). (**C-D**) Respective quantification of Caspase 3/7 activity in samples as in panels (A) and (B) respectively. n=4/ group, with each sample a pool of 6-10 embryos, Kruskal-Wallis (Dunn’s multiple comparison test). (**E**) Immunostaining of 4HNE (black) in NRP2 (red) lymphatic and CD31 (blue) blood dermal endothelial cells from the murine skin at embryonic day E13.5 and up to E16.5. Yellow boxes 1-3 appear in higher magnification on the right side of the panel with and without NRP2. Yellow dashed line: lymphatic vessel as marked from NRP2. Scale bars = 300 μm and 40 μm. (**F**) Oxidized to reduced ratio of MitoPerox levels in blood and lymphatic endothelial cells isolated from murine embryo skin on E15.5. n=4/ group, every sample is a pool of 4 embryos, Shapiro-Wilk normality test, unpaired Student’s t test. (**G**) Immunostaining of LYVE1 (red) lymphatic endothelial cells on postnatal (p) days 10, 16 and 22 in the murine cornea. Magnified panels show LYVE1 and 4HNE (black) levels of the black arrow pointed lymphatic vessels. Yellow dashed line: lymphatic vessel. Scale bars = 400 μm and 40 μm. (**H**) Oxidized to reduced ratio of MitoPerox levels in lymphatic endothelial cells isolated from postnatal murine corneas from p6 and up to p22. n=4/ group, every sample is a pool of 4 corneas, Kruskal-Wallis (Dunn’s multiple comparison test). (**I**) Oxidized to reduced ratio of MitoPerox levels in lymphatic endothelial cells isolated from p16 murine cornea after receiving treatment with solvent or 50 μmol/L Ferrostatin 1 (Fer1) from p14 and every 12 hours. n=4/ group, every sample is a pool of 4 embryos, Kolmogorov-Smirnov normality test, paired Student’s t test.

**Fig. S2.**
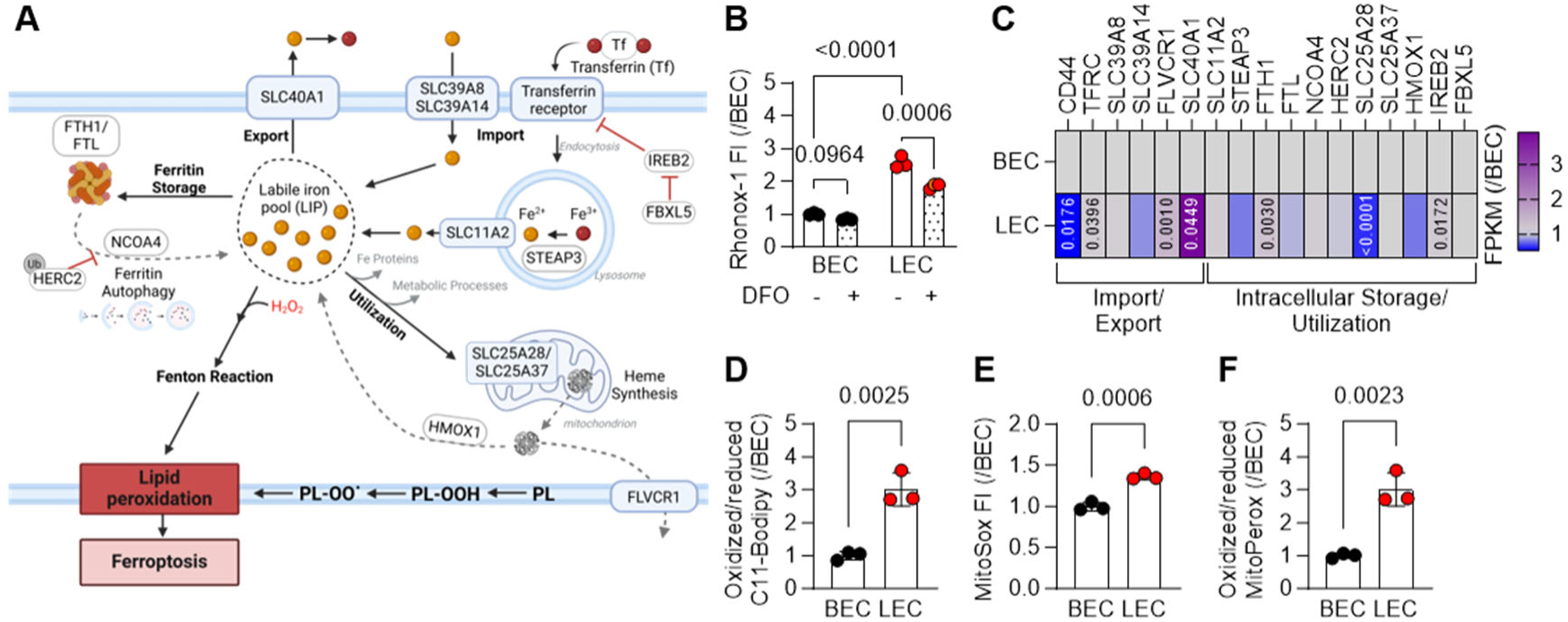
Increased labile iron accumulation correlates to increased lipid peroxidation in human blood but not lymphatic endothelial cells. (**A**) Schematic presentation of labile iron pool accumulation in the mammalian cells. (**B**) Fe^2+^ levels detected by the Rhonox-1 relative fluorescent intensity (FI) in human dermal lymphatic (LEC) and blood (BEC) endothelial cells exposed to solvent or the iron chelator deferoxamine (DFO, 100 μmol/L, 8 hours) following flow cytometry analysis. n=3/ group (in 3 technical replicates). Two way Anova, Fischer’s multiple comparison test. (**C**) Heatmap showing the relative transcriptional levels of the key iron homeostasis related mRNAs, in fragments per kilobase per million reads following RNA sequencing in human dermal lymphatic (LEC) and blood (BEC) endothelial cells. n=9/group. Kolmogorov-Smirnov normality test. Unpaired Student‘s t test. NCOA4: Nuclear receptor coactivator 4; FTH1: Ferritin heavy chain; FTL: Ferritin light chain; HERC2: E3 ubiquitin-protein ligase HERC2; FBXL5: F-box/LRR-repeat protein 5; SLC25A28: Mitoferrin-2; SLC25A37: Mitoferrin-1; HMOX1: Heme oxygenase 1; FLVCR1: Heme transporter; FLVCR1: Heme transporter FLVCR1; STEAP3: Metalloreductase STEAP3; SLC39A8: Metal cation symporter ZIP8; SLC39A14: Metal cation symporter ZIP14; SLC11A2: Natural resistance-associated macrophage protein 2; CD44: CD44 antigen; TFRC: Transferrin receptor protein 1; IREB2: Iron-responsive element-binding protein 2; SLC40A1: Solute carrier family 40 member 1. (**D**) Oxidized to reduced C11-BODIPY levels in human dermal lymphatic (LEC) and blood (BEC) endothelial cells following flow cytometry analysis. n=3/group (in 2 technical replicates). Sharipo-Wilk normality test. Unpaired Student‘s t test. (**E**) MitoSox fluorescent intensity (FI) in human dermal lymphatic (LEC) and blood (BEC) endothelial cells following flow cytometry analysis. n=3/group (in 2 technical replicates). Sharipo-Wilk normality test. Unpaired Student‘s t test. (**F**) Oxidized to reduced MitoPerox levels in human dermal lymphatic (LEC) and blood (BEC) endothelial cells following flow cytometry analysis. n=3/group (in 2 technical replicates). Sharipo-Wilk normality test. Unpaired Student‘s t test.

**Fig. S3.**
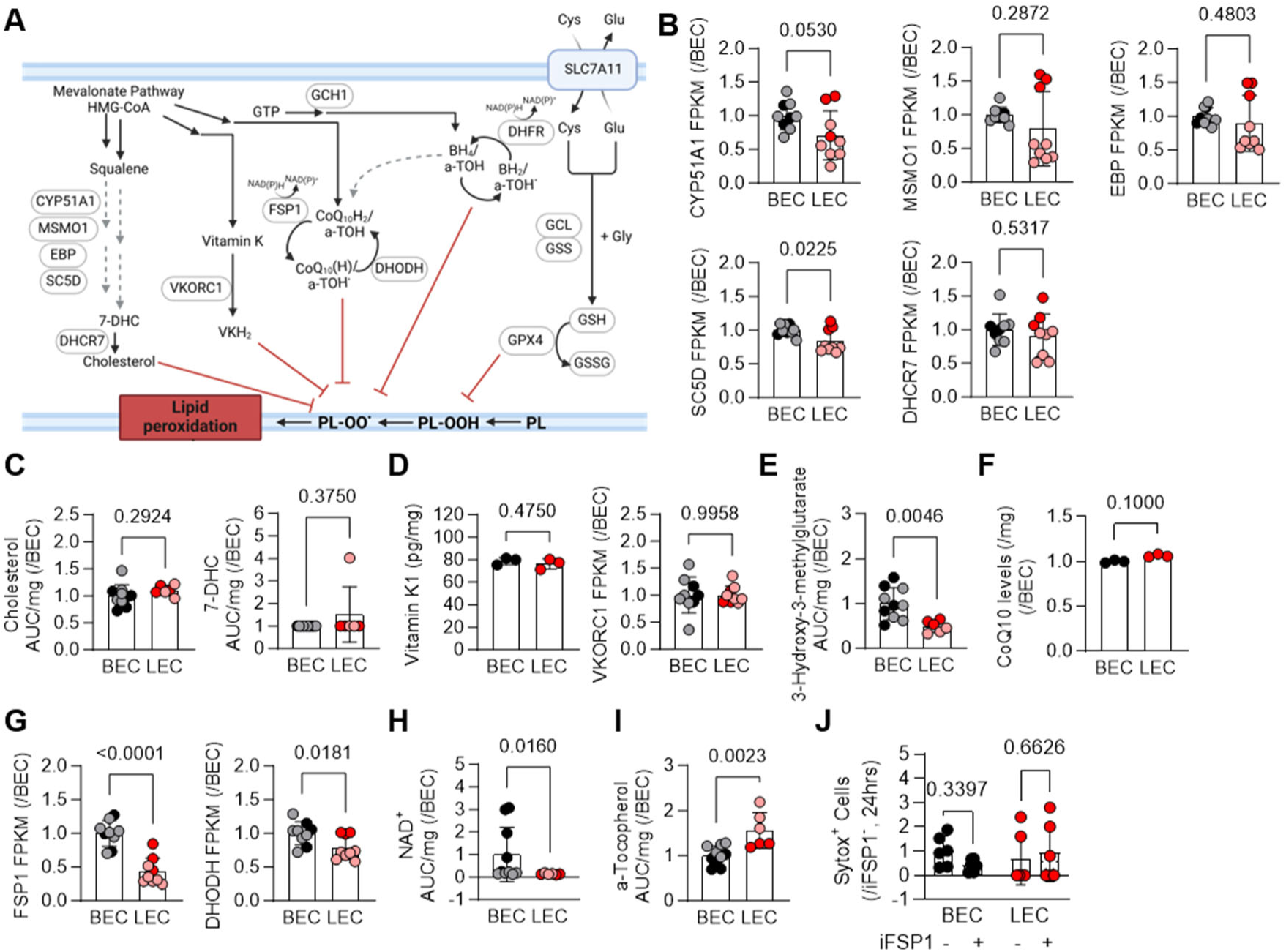
Lymphatic endothelial cells sensitivity to ferroptosis is independent from the mevalonate pathway. (**A**) Schematic showing the mevalonate ferroptotic suppressing pathways in mammalian cells. CYP51A1: Cytochrome P450 family 51 subfamily A member 1; MSMO1: Methylsterol monooxygenase 1; EBP: cholestenol delta-isomerase; SC5D: sterol-C5-desaturase; DHCR7: 7-dehydrocholesterol reductase; VCORC1: vitamin K epoxide reductase complex subunit 1; FSP1: Ferroptosis suppressor protein 1 ; GCH1: GTP cyclohydrolase 1; DHODH: dihydroorotate dehydrogenase ; DHFR: Dihydrofolate Reductase; GCL: glutamate-cysteine ligase: GSS: glutathione synthetase; GSH: Glutathione; GSSG: Glutathione disulfide; GPX4: glutathione peroxidase 4; SLC7A11: Solute carrier family 7 member 11; 7-DHC: 7-dehydrocholesterol; VKH2: Vitamin K hydroquinone; GTP: Guanosine Trisphosphate; CoQ10H2: Ubiquinol; a-TOH: a-Tocopherol; BH4: Tetrahydrobiopterin; Cys: Cysteine; Glu: Glutamate; Gly: Glycine; PL: Phospholipid. (**B**) Relative transcriptional levels of fragments per kilobase million of genes involved in the mevalonate to cholesterol biosynthesis in human lymphatic (LEC) and blood (BEC) endothelial cells. n=9/ group. Komogorov-Smirnov normality test, unpaired Student’s t-test. Lighter colors represent juvenile while darker colors adult cells. (**C**) Relative Cholesterol and 7-Dehydrocholesterol (7-DHC) levels in human lymphatic (LEC) and blood (BEC) endothelial cells. n=10/group. Kolmogorov-Smirnov normality test, unpaired Student’s t-test and Mann-Whitney test. Lighter colors represent proliferative and darker colors quiescent cells. (**D**) Relative Vitamin K1 levels and relative transcriptional levels of fragments per kilobase million of VKORC1 in human lymphatic (LEC) and blood (BEC) endothelial cells. Lighter colors represent proliferative or adult and darker colors quiescent or juvenile cells respectively. n=3/group and n=9/ group. Shapiro-Wilk normality test, unpaired Student’s t test. (**E**) Relative 3-Hydroxy-3-methylglutarate levels in human lymphatic (LEC) and blood (BEC) endothelial cells. n=10/group. Kolmogorov-Smirnov normality test, unpaired Student‘s t test. Lighter colors represent proliferative and darker colors quiescent cells. (**F**) CoQ10 levels in human lymphatic (LEC) and blood (BEC) endothelial cells. n=3/ group. Shapiro-Wilk normality test, unpaired Student’s t test. (**G**) Relative transcriptional levels of fragments per kilobase million of FSP1 and DHODH in human lymphatic (LEC) and blood (BEC) endothelial cells. n=9/ group. Kolmogorov-Smirnov normality test, unpaired Student’s t test. Lighter colors represent juvenile while darker colors adult cells. (**H**) Relative NAD^+^ levels in human lymphatic (LEC) and blood (BEC) endothelial cells. n=10/group. Kolmogorov-Smirnov normality test, Mann-Whitney test. Lighter colors represent proliferative and darker colors quiescent cells. (**I**) Relative a-Tocopherol levels in human lymphatic (LEC) and blood (BEC) endothelial cells. n=10/group. Kolmogorov-Smirnov normality test, unpaired Student’s t test. Lighter colors represent proliferative and darker colors quiescent cells. (**J**) Relative Sytox positive human lymphatic (LEC) and blood (BEC) endothelial cells treated with solvent or the FSP1 inhibitor (iFSP1, 3 μmol/L, 24 hours). n=6/group. Two way Anova, Fischer’s multiple comparison test.

**Fig. S4.**
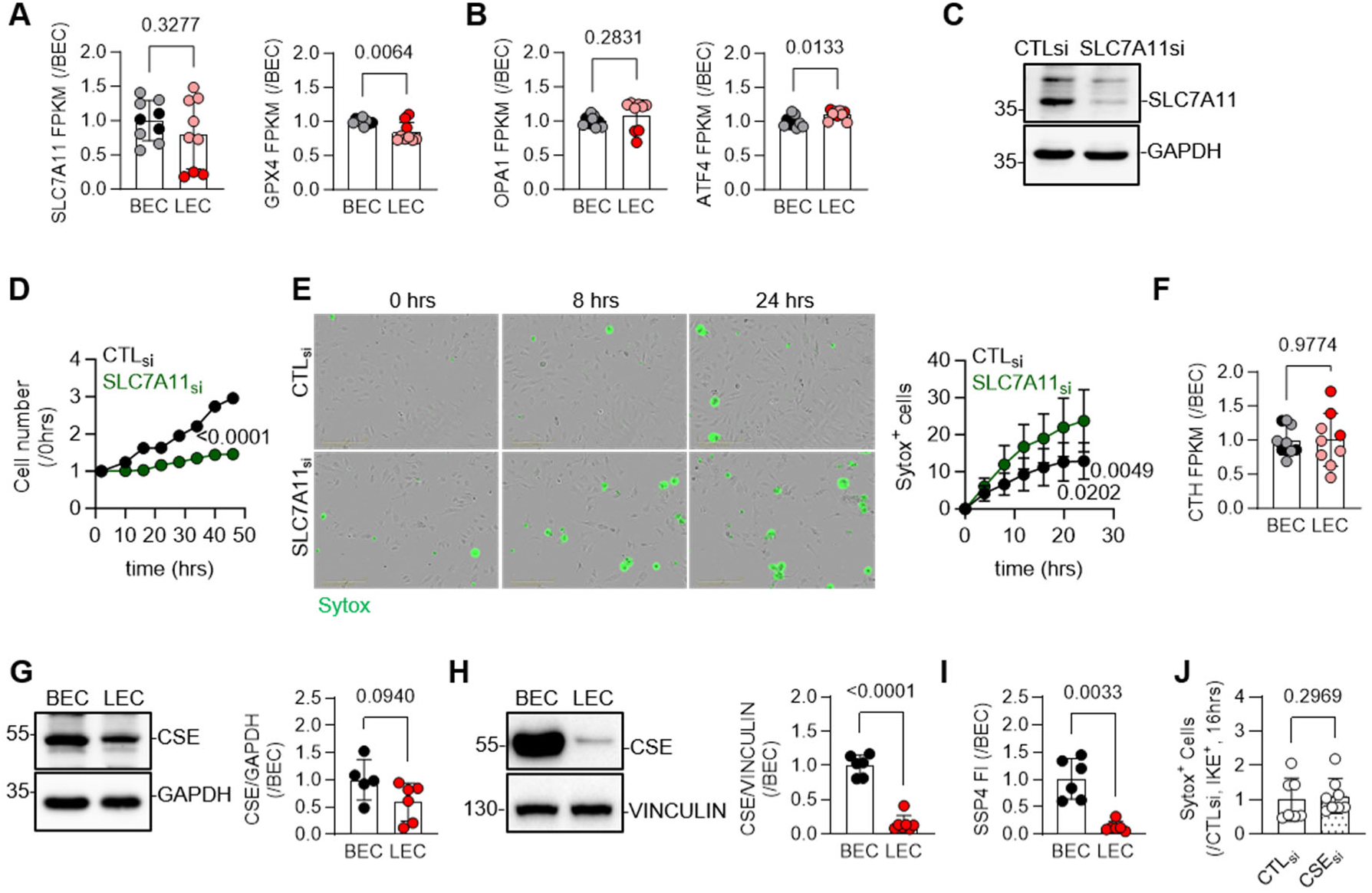
Lymphatic endothelial cells are sensitive to SLC7A11/CSE mediated ferroptosis. (**A-B**) Solute Carrier Family 7 Member 11 (SLC7A11), Glutathione Peroxidase 4 (GPX4) (A) and mitochondrial Dynamin-Like GTPase OPA1 (OPA1), Activating transcriptional factor 4 (ATF4) (**B**) transcriptional levels in human lymphatic (LEC) and blood (BEC) endothelial cells. n=9/ group. Kolmogorov-Smirnov normality test, unpaired Student’s t test. Lighter colors represent juvenile while darker colors adult cells. (**C**) Representative immunoblot of SLC7A11 and GAPDH in human dermal lymphatic (LEC) and blood (BEC) endothelial cells treated with siRNA targeting SLC7A11 (SLC7A11_si_) or a respective CTL (CTL_si_). Similar results were obtained in n=3 cell batches. (**D**) Relative cell number of human dermal lymphatic endothelial cells as in panel (C) for up to 48 hours. n=3/ group in 2 independent experiments. Two way Anova, Dunnet’s multiple comparisons test. (**E**) Representative images showing cell confuence and Sytox positive cells and respective quantification in samples as in panel D for up to 24 hours. n=3/ group in 3 independent experiments. Two way Anova, Dunnet’s multiple comparisons test. (**F**) CTH transcriptional levels in human dermal lymphatic (LEC) and blood (BEC) endothelial cells. n=9/ group. Kolmogorov-Smirnov normality test, unpaired Student’s t test. Lighter colors represent juvenile while darker colors adult cells. (**G**) Representative immunoblot of CSE and GAPDH in human dermal lymphatic (LEC) and blood (BEC) endothelial cells and respective quantification. n=5-6/ group. Kolmogorov-Smirnov normality test, unpaired Student’s t test. (**H**) Representative immunoblot of CSE and GAPDH in murine dermal lymphatic (LEC) and blood (BEC) endothelial cells isolated from the embryo skin on E15.5. and respective quantification. n=6/ group. Each is a pool of 4-6 embryos. Kolmogorov-Smirnov normality test, paired Student’s t test. (**I**) SSP4 fluorescent intensity in samples as in panel H. n=6/ group. Kolmogorov-Smirnov normality test, paired Student’s t test. (**J**) Sytox positive blood endothelial cells treated with a control siRNA (CTL_si_) or an siRNA directed against CSE (CSE_si_) for 72 hours and then treated with 10 µmol/L IKE for 16 hours. n=6-7/ group. Kolmogorov-Smirnov normality test, Wilcoxon test.

**Fig. S5.**
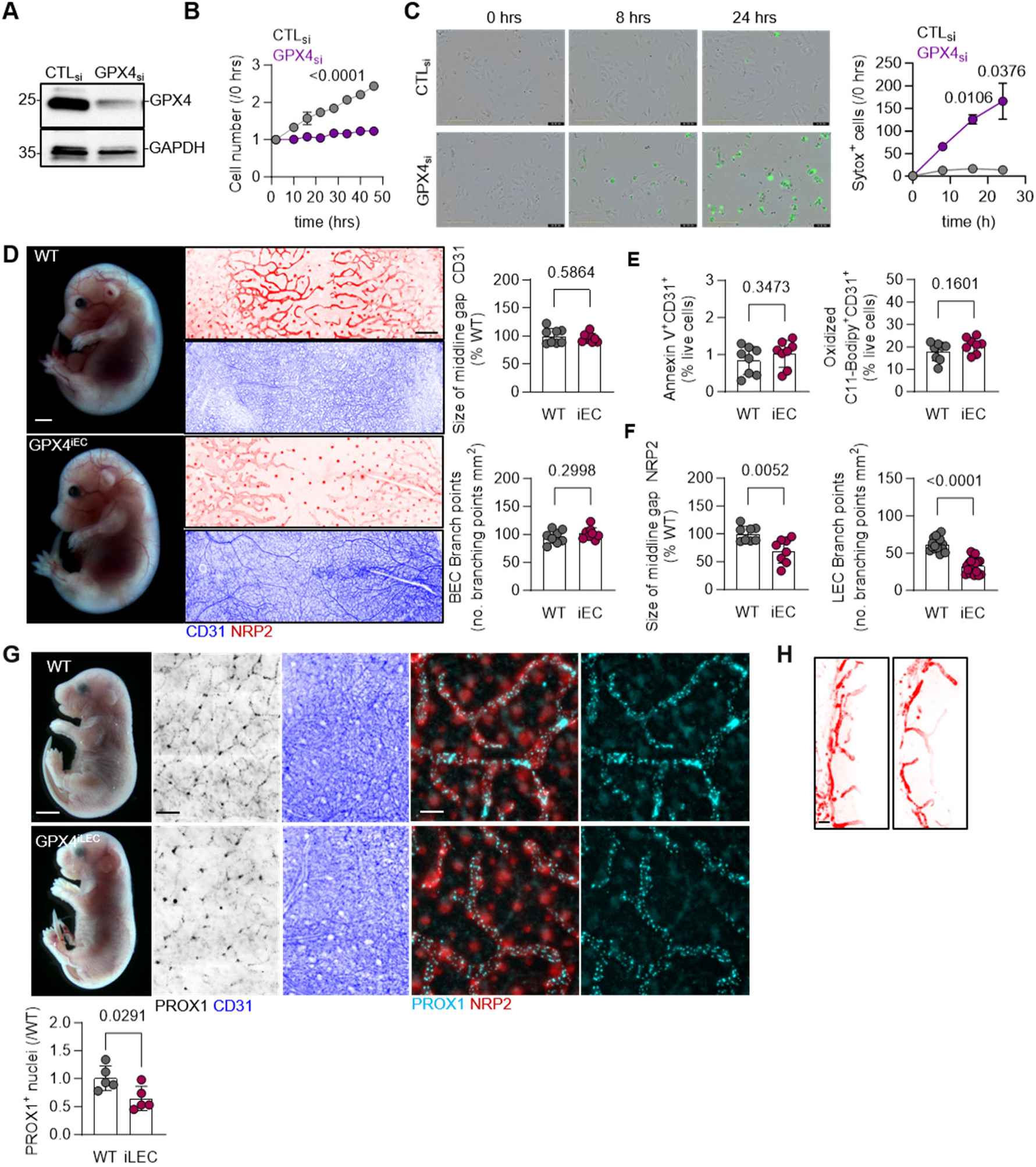
Loss of GPX4 does not impair blood endothelial cell growth. (**A**) Representative immunoblot of GPX4 and GAPDH in human lymphatic endothelial cells treated with a control siRNA (CTL_si_) or a siRNA directed against GPX4 (GPX4_si_). Similar results were obtained in 3 cell batches. (**B**) Relative cell number of lymphatic endothelial cells as in panel (A), for up to 48 hours. n=3/ group in 3 independent experiments. Two way Anova, Dunnet’s multiple comparisons test. (**C**) Representative images showing cell confluence and Sytox positive cells and respective relative quantification in samples as in panel (A), for up to 24 hours. n=3/ group in 3 independent experiments. Two way Anova, Dunnet’s multiple comparisons test. (**D**) Stereomicrographic images of E15.5 embryos and immunofluorescent images of CD31 (blue) blood and NRP2 (red) lymphatic dermal endothelial cells in Cdh5 Cre^-^ (WT) or Cre^+^ (iEC) mice treated with tamoxifen from E9.5-E12.5 to induce genetic deletion of endothelial cell *Gpx4*. Relative quantification of middline gap of CD31 positive vessels as % to WT and number of blood branch points in the same embryos. Scale bars= 1 mm and 200 μm. n=8 embryos/ group from 3 independent births. Shapiro-Wilk normality test, unpaired Student’s t test. (**E**) Percentage of double Annexin V^+^ or oxidized C11-Bodipy and CD31^+^ live cells in samples as in panel D. n=6-8 embryos/ group. Kolmogorov-Smirnov normality test, unpaired Student’s t test. (**F**) Relative quantification of middline gap of NRP2 positive vessels as % to WT and number of lymphatic branch points in the same embryos as in panel D. n=8 embryos/ group from 3 independent births. Kolmogorov-Smirnov normality test, unpaired Student’s t test. (**G**) Stereomicrographic images of E18.5 embryos and immunofluorescent images of CD31 (blue) blood and PROX1 (black) and zoom in of PROX1 (cyan) and NRP2 (Red) lymphatic dermal endothelial cells in PROX1 Cre^-^ (WT) or Cre^+^ (iLEC) mice treated with tamoxifen from E14.5-E16.5 to induce genetic deletion of blood endothelial cell *Gpx4*. Relative levels of PROX1 positive nuclei in the same samples. Scale bars = 1 mm, 200 μm and 50 μm. n=5 embryos/ group from 2 independent births. Kolmogorov-Smirnov normality test, unpaired Student’s t test. (**H**) Immunostaining of LYVE1 (red) on postnatal day (p) 22 murine cornea after inducing lymphatic specific deletion of GPX4 (iLEC) from p16 to p18, and respective quantification of lymphatic vessel length. n=4 corneas/ group. Scale bars = 100 μm. Kolmogorov-Smirnov normality test, unpaired Student’s t test.

**Fig. S6.**
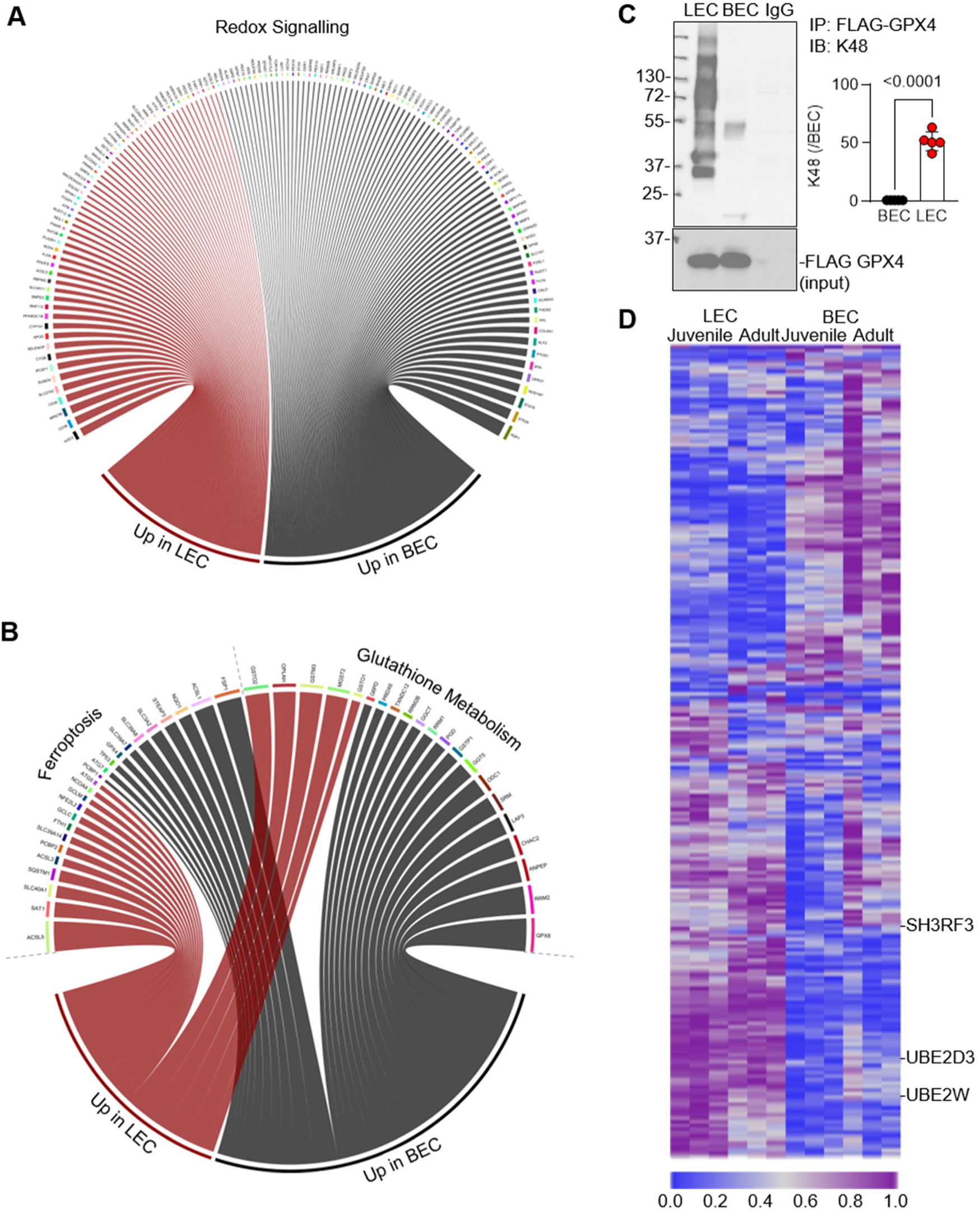
Lymphatic and blood endothelial cells exhibit differential antioxidant and protein degradation transcriptional signatures. (**A-B**) Circos plot showing differential expression of Redox signaling (merge from KEGG: hsa04146, Gene Ontology: GO0051341, GO0006979, Gene set enrichment analysis (GSEA): M39645, M5938), Ferroptosis (merge from KEGG: hsa04216, Gene Ontology: GO0097707) and Glutathione metabolism (merge from: KEGG: hsa00480, Gene set enrichment analysis (GSEA): M39388) transcripts following RNA sequencing of blood (BEC) and lymphatic (LEC) endothelial cells. Differentially expressed genes have log₂ fold change (|log₂FC| > 0.5) and an adjusted p-value (padj ≤ 0.05). n=6/ group. (**C**) Representative immunoblot for K48 polyubiquitination from immunoprecipitated 3xFlag-GPX4 in human blood (BEC) and lymphatic (LEC) endothelial cells. n=6/ group. Kolmogorov-Smirnov normality test. Unpaired Student‘s t test. (**D**) Heatmap showing ubiquitin ligase and deubiquitinase mRNA transcripts in samples as in panel A. n=6/ group.

**Fig. S7.**
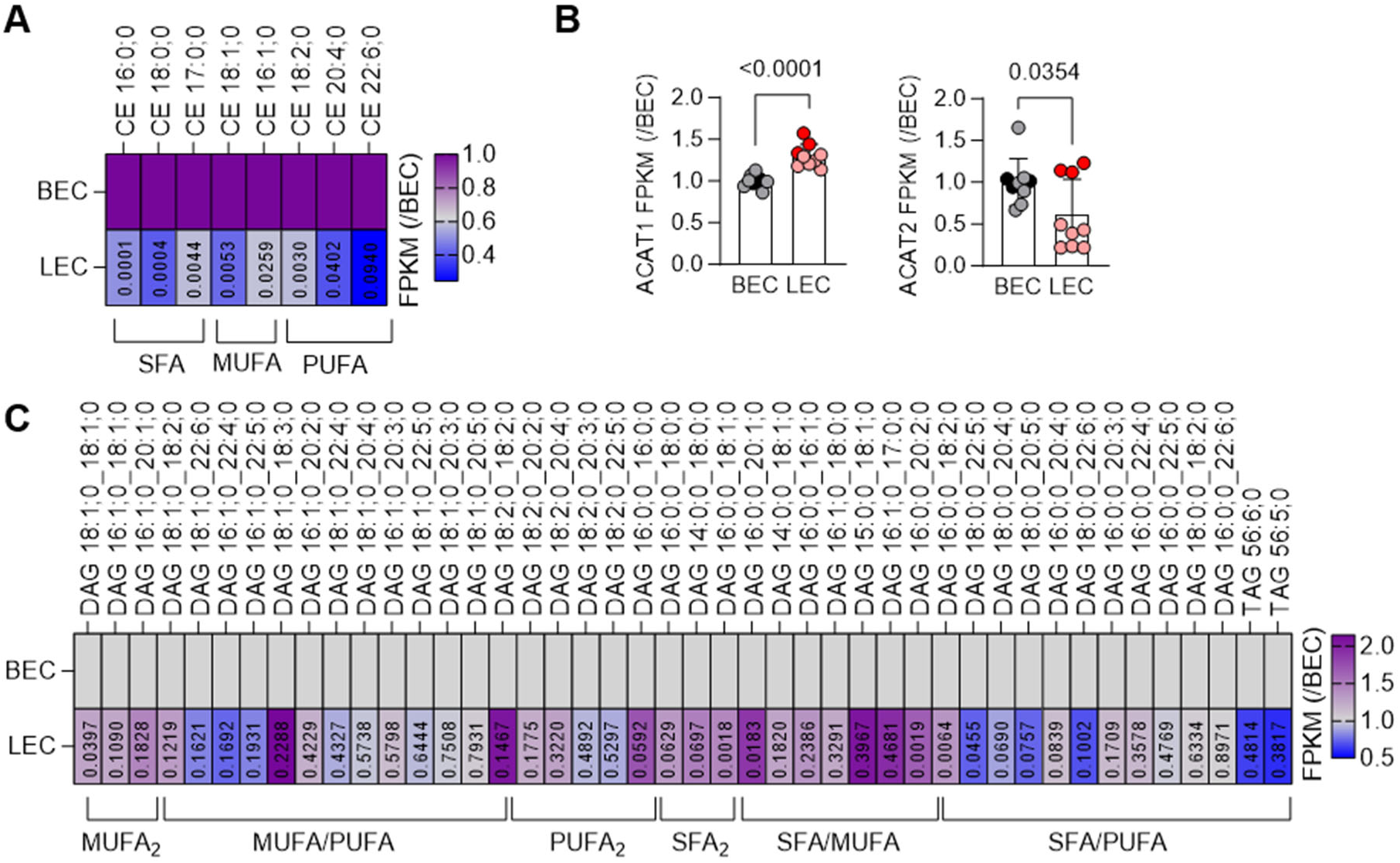
Lipidomic signature of cholesterol esterified fatty acids, diacyl and triacyl glycerolipids. (**A**) Heatmap showing relative levels of cholesterol ester fatty acids in human dermal blood (BEC) and lymphatic (LEC) endothelial cells. n= 3/ group. Shapiro-Wilk normality test, unpaired Student’s t test. (**B**) Relative transcriptional levels of fragments per kilobase million of the cholesterol ester synthetizing enzymes Acetyl-Coenzyme A acetyltransferase 1 and 2 (ACAT1/ 2). n=9/group. Kolmogorov-Smirnov normality test. Unpaired Student‘s t test. Light colors represent juvenile, while darker colors adult endothelial cells. (**C**) Heatmap showing relative levels of diacylglycerol and triacyl glycerol fatty acids in human dermal blood (BEC) and lymphatic (LEC) endothelial cells. n= 3/ group. Shapiro-Wilk normality test, unpaired Student’s t test.

**Fig. S8.**
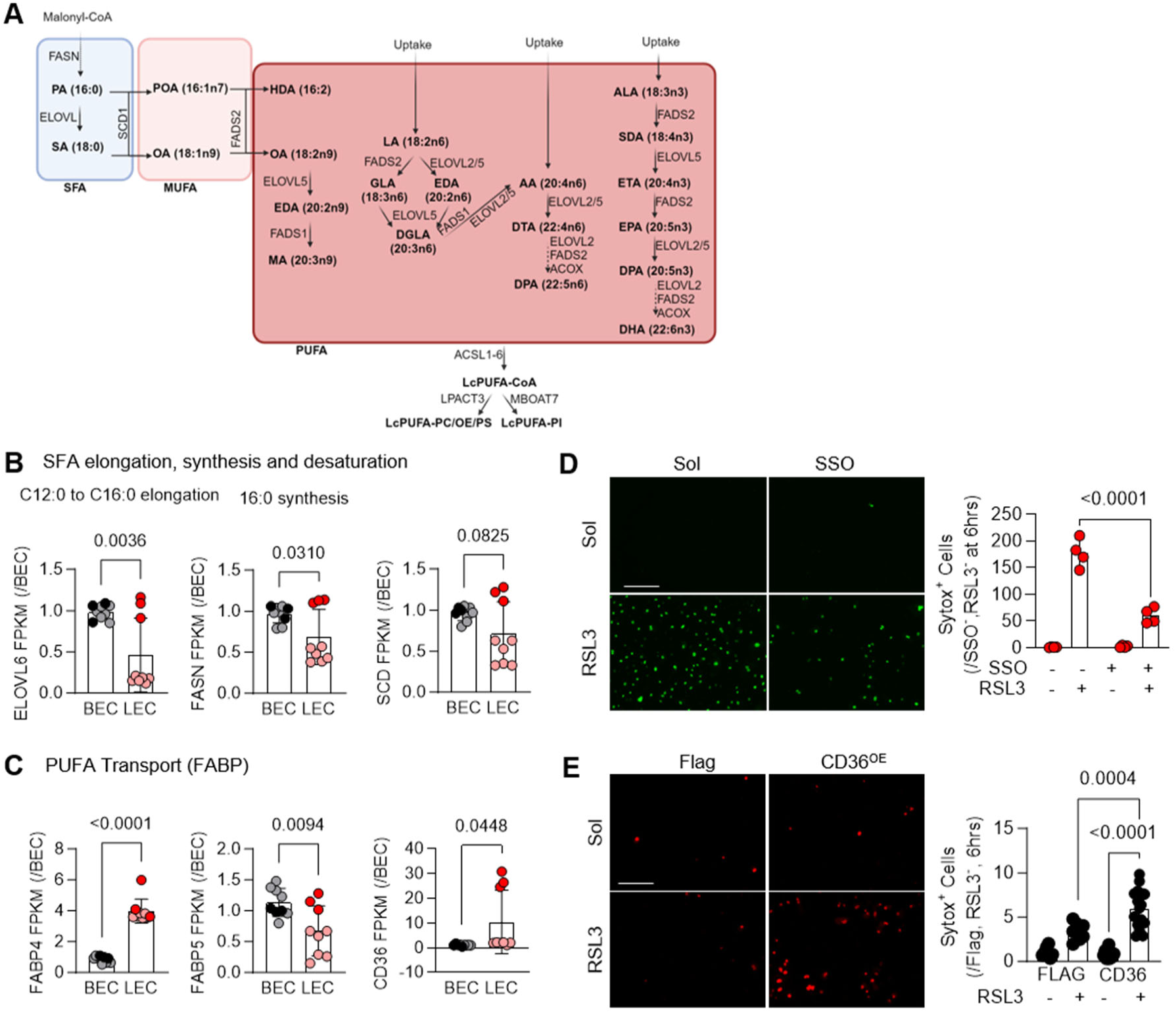
PUFA transport differs between blood and lymphatic endothelial cells. (**A**) Schematic showing SFA, MUFA and PUFA utilization and biosynthesis in mammalian cells. (**B**) Relative fragments per killobase million of the SFA C12:0 to C16:0 elongation (ELOVL6: Very long chain fatty acid elongase), synthesis (FASN: Fatty acid synthase) and desaturation (SCD1: Stearoyl-CoA Desaturase) genes following RNA sequencing in human blood (BEC) and lymphatic (LEC) endotheliall cells. n=9/group. Kolmogorov-Smirnov normality tests, unpaired Student‘s t test. Light colours represent juvenile while dark colours adult cells. (**C**) Relative fragments per killobase million of the PUFA transporting genes (FABP4: Fatty acid binding protein 4; FABP5, Fatty acid binding protein 5; CD36: Platelet glycoprotein 4) in samples as in panel (B). n=9/group. Kolmogorov-Smirnov normality test, unpaired Student‘s t test. Light colours represent juvenile while dark colours adult cells. (**D**) Relative Sytox positive human dermal lymphatic endothelial cells, cultured in the presence or absence of 50 µmol/L Sulfosuccinimidyl oleate (SSO, 24 hours) and subsequently exposed to RSL3 (5 μmol/L, 6 hrs), n=4/ group. Two way Anova, Fischer’s multiple comparison test. (**E**) Relative Sytox positive human blood (BEC) endothelial cells infected with a Flag lentivirus or a virus overexpressing the human CD36, and subsequently exposed to RSL3 (5 μmol/L, 6 hours). n=6-9/ group. Two way Anova, Fischer’s multiple comparison test.

**Fig. S9.**
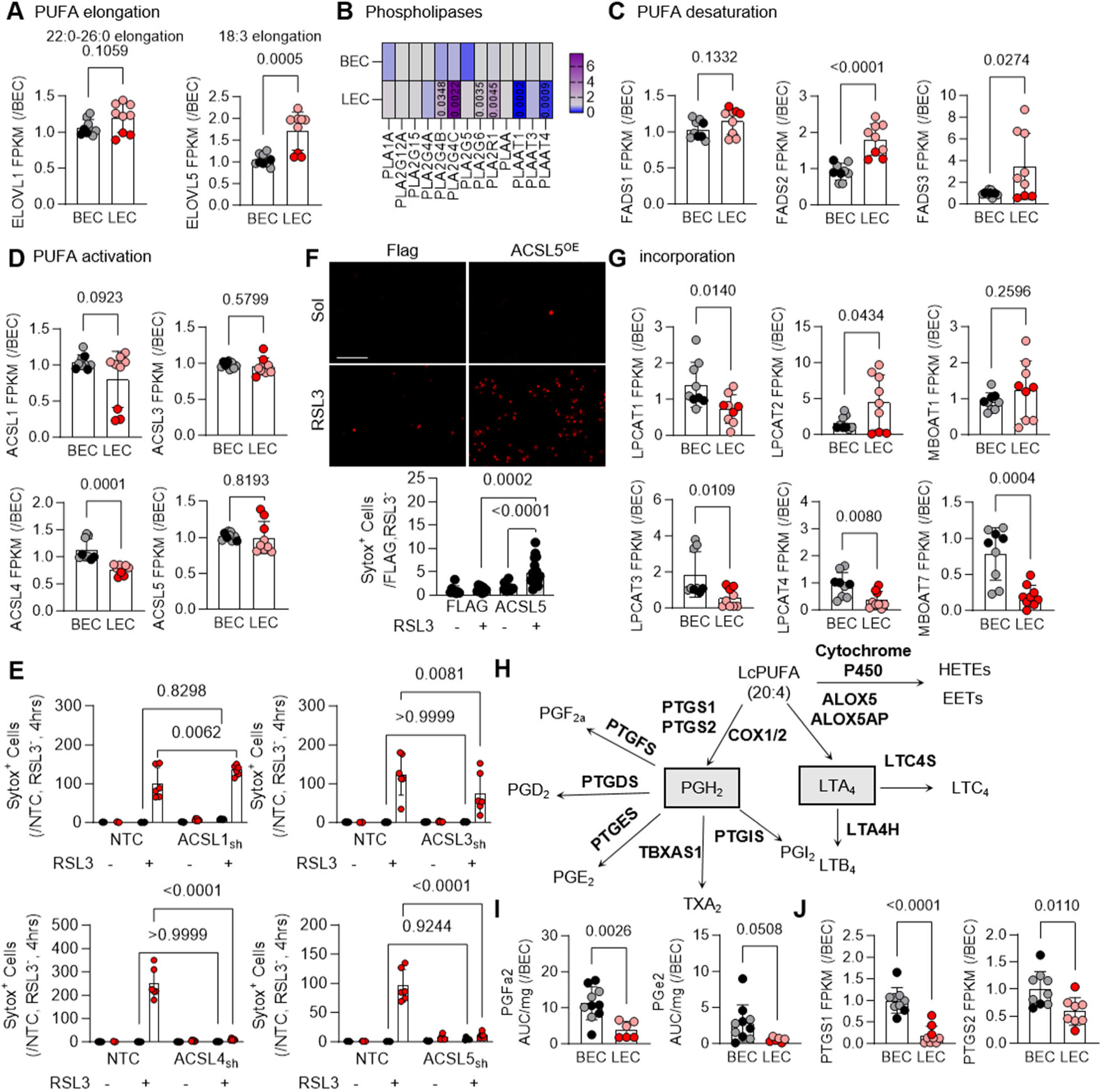
Blood endothelial cells activate less PUFAs and utilize them to generate prostaglandins. (**A**) Relative transcriptional levels of fragments per kilobase million of the PUFA elongation (ELOVL1, 5: Very long chain fatty acid elongase 1,5) genes following RNA sequencing in human blood (BEC) and lymphatic (LEC) endothelial cells. n=9/group. Kolmogorov-Smirnov normality test, unpaired Student‘s t test. Light colors represent juvenile while dark colors adult cells. (**B**) Heatmap showing relative transcriptional levels of fragments per kilobase million of phospholipase levels (PLAA: Phospholipase A-2-activating protein; PLA2R1: Secretory phospholipase A2 receptor; PLAAT3: Phospholipase A and acyltransferase 3; PLA2G4C: Cytosolic phospholipase A2 gamma; PLA2G4A: Cytosolic phospholipase A2; PLA2G12A: Group XIIA secretory phospholipase A2; PLA2G15: Phospholipase A2 group XV) in samples as in panel (A). n=9/group. Kolmogorov-Smirnov normality test, unpaired Student‘s t-test. (**C**) Relative transcriptional levels of fragments per kilobase million of the PUFA desaturation (FADS1: Acyl-CoA (8–3)-desaturase; FADS2: Acyl-CoA 6-desaturase; FADS3: Fatty acid desaturase) genes in samples as in panel (A). n=9/group. Kolmogorov-Smirnov normality test, unpaired Student‘s t test. Light colors represent juvenile while dark colors adult cells. (**D**) Relative transcriptional levels of fragments per kilobase million of the PUFA activation (ACSL1,3,4,5: Acyl-CoA synthetase long-chain family member 1,3,4,5) genes in samples as in panel (A). n=9/group. Kolmogorov-Smirnov normality test, unpaired Student‘s t test. Light colors represent juvenile while dark colors adult cells. (**E**) Relative Sytox positive human lymphatic endothelial cells infected with a non targeting control CRISPR-Cas9 lentivirus or viruses against the human ACLS 1,3,4,5 and subsequently exposed to RSL3 (5 μmol/L, 6 hrs). n=6-9/ group. Two way Anova, Fischer’s multiple comparison test. (**F**) Relative Sytox positive human blood endothelial cells infected with a Flag lentivirus or a virus overexpressing the human ACLS5, and subsequently exposed to RSL3 (5 μmol/L, 6 hrs). n=6-9/ group. Two way Anova, Fischer’s multiple comparison test. (**G**) Relative transcriptional levels of fragments per kilobase million of the PUFA incoroporation into the membranes (LPCAT1,2,3,4: Lysophosphatidylcholine acyltransferase and MBOAT1,7: Lysophospholipid acyltransferase 1, 7) genes in samples as in panel (A). n=9/group. Kolmogorov-Smirnov normality test, unpaired Student‘s t test. Light colors represent juvenile while dark colors adult cells. (**H**) Schematic presentation of PUFA derived bioactive lipids in mammalian cells. ALOX5: Arachidonate 5-Lipoxygenase; ALOX5AP: Arachidonate 5-lipoxygenase-activating protein; LTC4S: Leukotriene C4 Synthase; LTA4H: Leukotriene A4 Hydrolase; COX1/2: Cyclooxygenase 1/2; PTGS1: Prostaglandin-Endoperoxide Synthase 1; PTGS2: Prostaglandin-Endoperoxide Synthase 2; PTGFS: Prostaglandin D2 Synthase; PTGDS: Prostaglandin D2 Synthase; PTGES: Prostaglandin E Synthase; TBXAS1: Thromboxane A Synthase 1; PTGIS: Prostaglandin I2 Synthase; HETEs: Hydroxyeicosatetraenoic acids; EETs: Epoxyicosatrienoic Acid; LTC_4_: Leukotriene C4; LTB_4_: Leukotriene B4; LTA_4_: Leukotriene A4; PGI_2_: Prostacyclin; TXA_2_: Thromboxan A2; PGE_2_: Prostaglandin E2; PGD_2_: Prostaglandin D2; PGF_2a_: Prostaglandin F2a; PGH_2_: Prostaglandin H2. (**I**) Relative levels of PGE2 and PGF2a in samples as in panel A. n=6-7/group. Kolmogorov-Smirnov normality test, unpaired Student‘s t test. (**J**) Relative transcriptional levels of fragments per kilobase million of the genes responsible for the arachidonic acid conversion to prostaglandins (PTGS1, 2) in samples as in panel (A). n=9/group. Kolmogorov-Smirnov normality test, unpaired Student‘s t test.

**Fig. S10.**
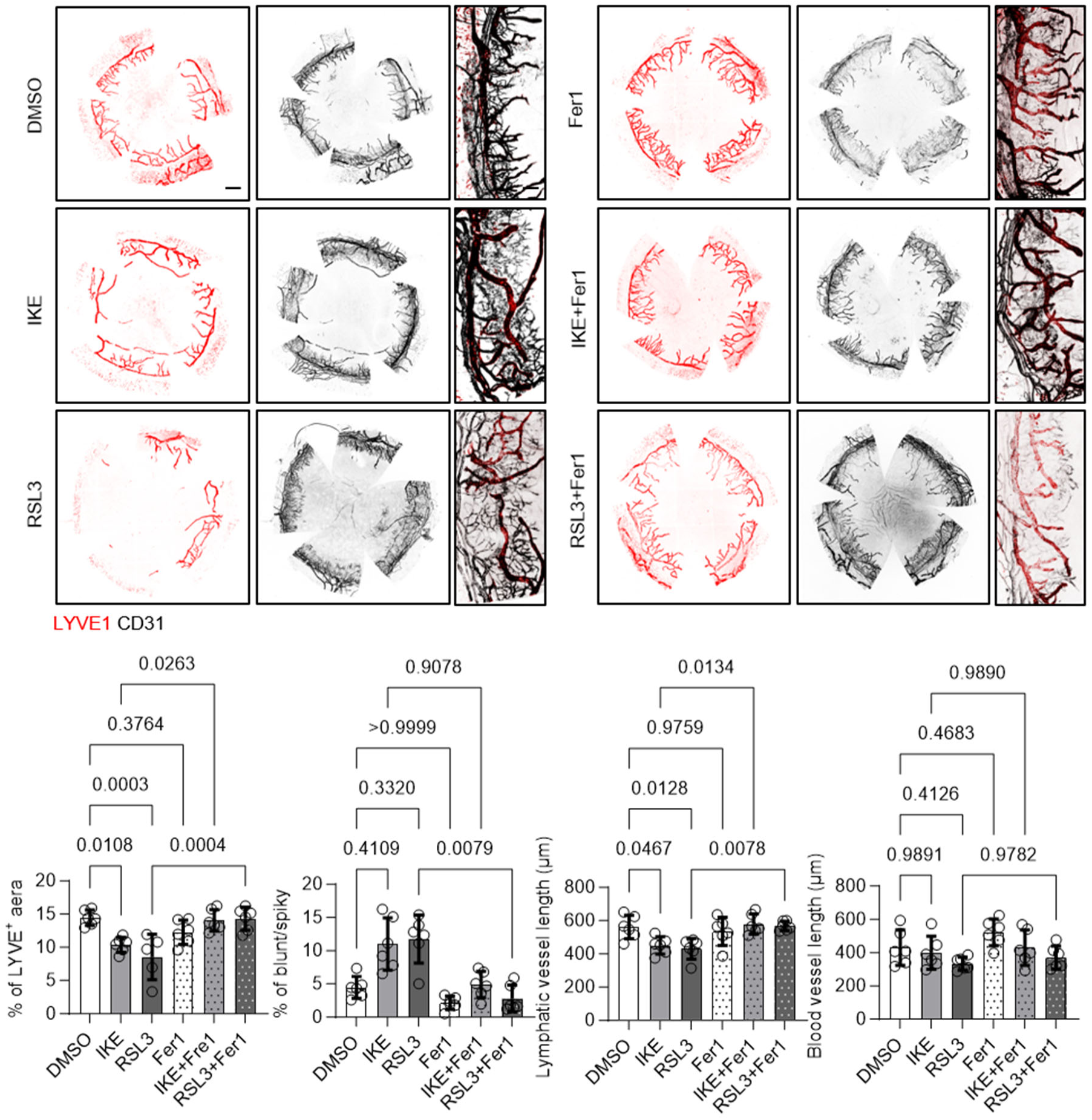
Pharmacological induction of ferroptosis impacts on lymphatic growth after injury. Immunostaining of LYVE1 (red) lymphatic and CD31 (black) blood endothelial cells in adult corneas from wild type mice treated with DMSO, IKE (10 μmol/L), RSL3 (10 μmol/L) with or without the addition of Ferrostatin 1 (Fer1, 50 μmol/L) following alkali burn cornea injury for 7 days. Scale bars = 500 µm and 100 µm. n= 5-6/ group. Kolmogorov-Smirnov normality test. One way Anova (Tukey‘s multiple comparison test).

## Notes

### Competing Interest Statement

The authors have declared no competing interest.

